# Multi-omics investigation reveals molecular determinants of cancer cell evolution on soft extracellular matrix

**DOI:** 10.64898/2025.12.17.694784

**Authors:** Sarthak Sahoo, Sejal Khanna, Ting-Ching Wang, Tanmay P. Lele, Mohit Kumar Jolly

## Abstract

Cancer cell adaptation to their physical tumor microenvironment is a key driver of malignancy. Recent experimental evolution experiments show that the soft extracellular matrix (ECM) can impose a selection pressure on genetically variable tumor populations. Over months of sustained culture, the selection pressure leads to enrichment of specific genetic variants with high fitness, but the mechanisms underlying the high fitness of these soft-selected clones are not fully understood. Here, we used a combination of RNA-seq, ATAC-seq, and RRBS-seq to compare soft-selected populations with non-selected ancestral populations cultured on soft ECM. We demonstrate that ancestral populations grown on soft ECM for short durations are characterized by a stressed cell state with low fitness marked by cell cycle arrest and distinct metabolic shifts, whereas sustained culture selects for a robust proliferative phenotype. Mechanistically, selected cells exhibit a silenced ancestral stress response through epigenetic modifications, characterized by reduced chromatin accessibility and de novo DNA methylation, including CDH1 promoter hypermethylation. This repressive landscape supports a high-fitness state defined by elevated MYBL2 and FAK levels. An *in-silico* mechanism-based model shows that these molecular differences, together with high YAP1 nuclear localization in soft-selected cells, are salient features of genetic clones capable of FAK upregulation. These findings uncover a coordinated genetic and epigenetic mechanism driving cancer cell evolution in mechanically soft microenvironments.

## Introduction

Cancer development and progression depend on the ability of cells to sense and adapt to the physical properties of their surroundings. Among these properties, the stiffness of the extracellular matrix (ECM) plays a pivotal role in shaping malignant behavior. In many solid tumors, such as breast cancer, increased tissue stiffness is not merely a passive consequence of disease but an active determinant of tumor initiation and progression (1–4). Breast cancer cells must continually adapt to stiffness variations within the primary tumor during disease progression (5–7) and to softer microenvironments during dissemination and colonization at distant sites such as the brain and bone marrow (6–8). Adaptation to ECM stiffness helps cells to maintain proliferative signaling, resist apoptosis, and acquire invasive and metastatic capabilities (7,8), making ECM stiffness a central regulator of cancer cell plasticity and aggressiveness.

Cells perceive and translate physical cues into biochemical signals through a process known as mechanotransduction. This response is initiated at focal adhesions, where transmembrane integrin receptors link the ECM to the intracellular actin cytoskeleton. In a stiff environment, this physical linkage generates high intracellular tension, activating signaling hubs such as Focal Adhesion Kinase (FAK) and the GTPase RHOA (9). This cascade culminates in the activation and nuclear translocation of the transcriptional co-activator YAP, a master regulator of the mechanical response (10). Once in the nucleus, YAP partners with TEAD transcription factors to orchestrate a program that enhances proliferation (11,12) and provides a survival advantage.

While these pathways explain the cell’s plastic response to ECM stiffness, their role in cancer evolution, a characteristic of cancer, is less understood. Cancer evolution involves microenvironmental selection of fit genetic variants from a population of cells with genetic variation. We recently showed that ECM stiffness can exert selection pressure on MDA-MB-231 breast cancer cells (13). Prolonged culture of a genetically variable population of cells on soft ECM hydrogels over months resulted in the enrichment of a few genetic variants that displayed unusual properties, including high cell spreading, YAP nuclear localization, and YAP-dependent proliferation. Clonal populations cultured for several months on soft ECM did not display such behaviors. The mechanisms that underlie the differential fitness of the soft-selected cells are not well understood.

Here, we combined multi-omics profiling (RNA-seq, ATAC-seq, and RRBS-seq) and computational modeling to understand the differences between soft-selected cells cultured on soft ECM for several months and ancestral cells cultured on soft ECM for days. Our results demonstrate that both epigenetic and signaling mechanisms account for the fitness of the soft-selected genetic variants.

## Results

### WGCNA Identifies Distinct Transcriptional Programs Driving Phenotypic Differences During Selection on Soft and Stiff Substrates

To elucidate selection driven transcriptional differences underpinning the observed evolution of cancer cells on soft and stiff substrates, we investigated previously performed bulk RNA-sequencing (RNA-seq) of MDA-MB-231 cells (GSE255829); (13)). We analyzed four distinct experimental groups, each with four replicates: cells from the ancestral population cultured short-term on soft (Soft Ancestral) or stiff (Stiff Ancestral) substrates, and cells evolutionarily selected for 75 days (approximately 40 cell generations) on soft (Soft Selected) or stiff (Stiff Selected) substrates (**Fig. 1A**). Phenotypically, the Soft Ancestral cells are characterized by a lower net growth rate along with a distinctive lack of nuclear YAP1 levels. The YAP1 localization as well as growth rates are tandemly enriched once these cells are selected on soft substrates. Such drastic changes are not seen in the context of the Stiff Selected cells when compared to Stiff Ancestral cells (13).

**Figure 1.**
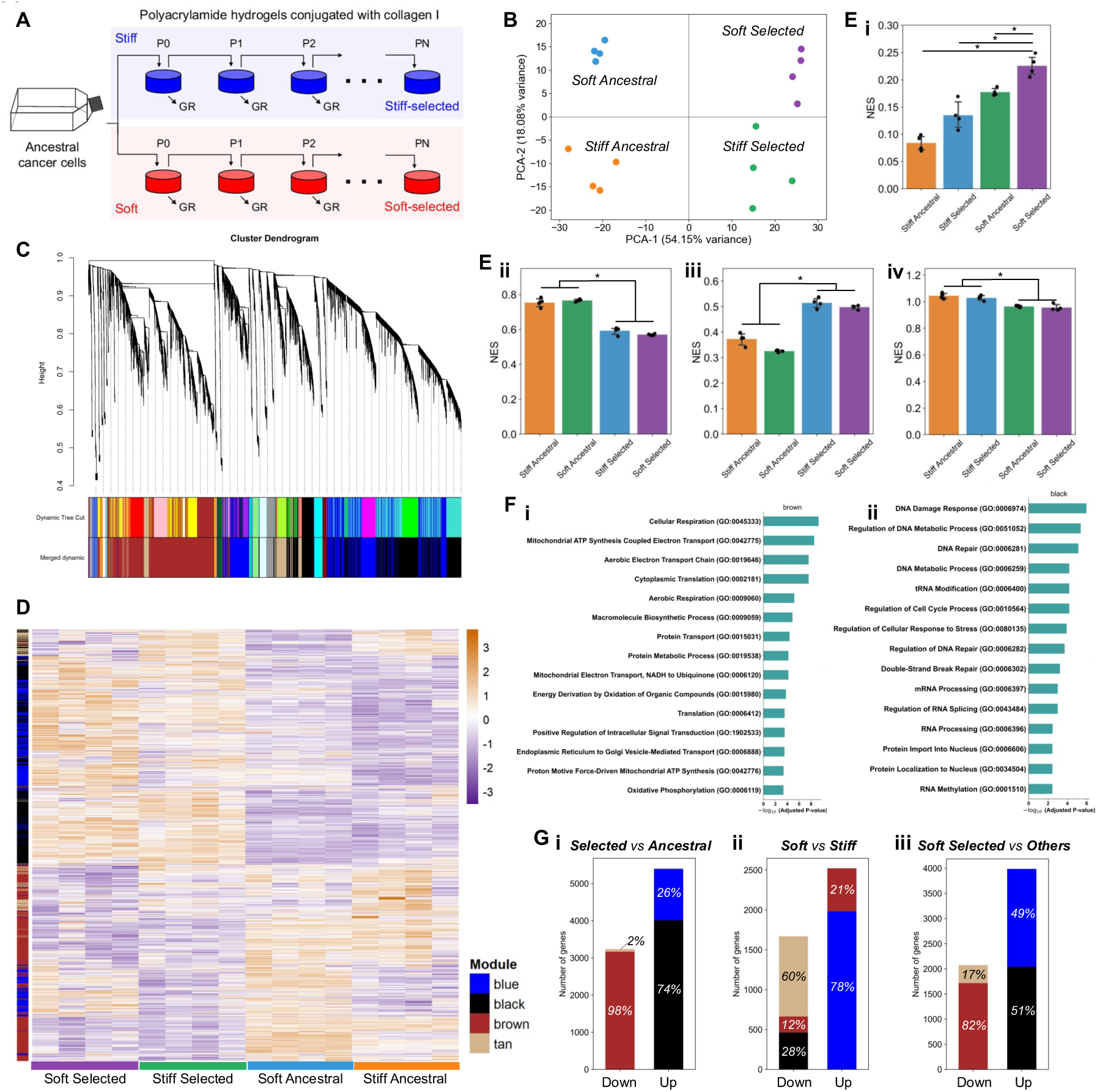
WGCNA Identifies Transcriptional Programs for Soft Substrate Phenotypic Differences. **(A)** Schematic of the experimental evolution setup, figure taken from reference (13). **(B)** PCA plot of RNA-seq samples. PC1 (54.15% variance) separates Selected vs. Ancestral groups. PC2 (18.08% variance) separates Soft vs. Stiff conditions. **(C)** WGCNA gene clustering dendrogram showing four distinct co-expression modules (blue, tan, black, brown) identified by topological overlap. **(D)** Expression heatmap of WGCNA module genes (rows) across all replicates and conditions (columns). Side-colors denote modules from (C). Orange = high expression, purple = low expression (z-normalized). **(E)** Module activity quantified by ssGSEA. Bar plots show mean scores (± standard deviation) for the blue (soft-selected), brown (ancestral), black (selected), and tan (stiff) modules. p < 0.05 (Student’s t-test). **(F)** Gene Ontology (GO) enrichment analysis of top biological processes for the (i) brown “ancestral” module (e.g., aerobic respiration) and (ii) black “selected” module (e.g., DNA repair, protein import to nucleus). **(G)** Contribution of WGCNA modules to differentially expressed gene (DEG) sets. Stacked bar plots show the module composition (in percent) of up-regulated (Up) and down-regulated (Down) genes for three key comparisons: (i) Selected vs. Ancestral, (ii) Soft vs. Stiff, and (iii) Soft Selected vs. Others.

Principal Component Analysis (PCA) of corresponding transcriptomic data revealed clear clustering by experimental condition (**Fig. 1B**). The primary axis of variation, Principal Component 1 (PC1), accounted for 54.15% of the variance and clearly separated the selected populations (Soft Selected and Stiff Selected) from the ancestral populations (Soft Ancestral and Stiff Ancestral). This indicates that the long-term selection process itself, irrespective of stiffness, was the dominant driver of transcriptional changes. The second axis, Principal Component 2 (PC2), explained 18.08% of the variance and distinguished the samples based on substrate stiffness, separating soft (Soft Selected and Soft Ancestral) from stiff (Stiff Selected and Stiff Ancestral) conditions (**Fig. 1B**).

To move beyond simple differential expressions across the different experimental groups and identify robust modules of co-expressed genes associated with these experimental conditions, we employed Weighted Gene Co-expression Network Analysis (WGCNA) (14). This unbiased approach can analyze large, potentially noisy lists of differentially expressed genes. It identified four distinct gene modules (**Fig. 1C**) labeled as blue, tan, black, and brown. A heatmap of the genes within these modules illustrates their distinct expression patterns across the four conditions, indicating specific expression profiles that associate with the four conditions (**Fig. 1D**).

We quantified the activity of each module using single-sample Gene Set Enrichment Analysis (ssGSEA) to define their functional identity (**Fig. 1E**). The blue module was identified as a “soft-selected” program (**Fig. 1E, i**). Its score was highest in the Soft Selected condition, significantly higher than all other conditions (Student’s t-test, p < 0.05). The Soft Ancestral condition also showed a higher score for this module compared to stiff samples, suggesting the blue module also comprises genes that belong to the “soft” program. The black module was assigned to a general “selected” program, as it was significantly elevated in both selected conditions (Soft Selected and Stiff Selected) compared to their ancestral counterparts (**Fig. 1E, ii**). The brown module was defined as an “ancestral” program, with significantly higher expression in both ancestral groups (Soft Ancestral and Stiff Ancestral) relative to the selected conditions (**Fig. 1E, iii**). The tan module emerged as a “stiff” program, exhibiting the highest scores in the stiff-cultured samples (Stiff Ancestral and Stiff Selected) regardless of selection status (**Fig. 1E, iv**).

We then performed Gene Ontology (GO) analysis of the modules representing the main axis of variability (ancestral vs. selected) to reveal their biological functions (**Fig. 1F**). The brown “ancestral” module was highly enriched for genes involved in metabolic pathways, particularly aerobic respiration and oxidative phosphorylation, suggesting the ancestral cells exist in a metabolically active basal state (**Fig. 1F, i**). In contrast, the black “selected” module was enriched for processes related to DNA repair, transcription, and protein import into the nucleus, pointing to a selected state focused on genomic maintenance and robust transcriptional activity associated with proliferation (**Fig. 1F, ii**).

Finally, we assessed how these modules contribute to the sets of differentially expressed genes (DEGs) in key comparisons (**Fig. 1G**). When comparing Selected vs Ancestral conditions taken together (**Fig. 1G, i**), the up-regulated genes were overwhelmingly dominated by the black “selected” program (74%) and the blue “soft-selected” program (26%). Conversely, down-regulated genes almost exclusively belonged to the brown “ancestral” program (98%), confirming the transcriptional shift away from the ancestral state. The comparison between soft vs stiff conditions taken together (**Fig. 1G, ii**) showed that up-regulated genes (higher in soft) were predominantly from the blue “soft-selected” program (78%). Down-regulated genes (higher in stiff) were strongly dominated by the tan “stiff” program (60%), with smaller contributions from the black (28%) and brown (12%) modules. To isolate the unique signature of phenotypic shift to the soft environment, we compared Soft Selected replicates to the other conditions (**Fig. 1G, iii**). The up-regulated gene set defining this state was an almost equal mix of the black “selected” program (51%) and the blue “soft-selected” program (49%). This demonstrates that the resulting phenotype is a composite of both general selection-enriched and soft substrate specific gene programs. As expected, the most strongly down-regulated genes were from brown “ancestral” program (82%).

Taken together, this transcriptomic analysis reveals that long-term culture selects programs involved in proliferation and DNA maintenance from a parental cell population enriched in metabolically active gene expression programs.

### Pathway-Level Analysis Reveals an Enrichment of a Low Fitness Mechanosensitive State Prior to an Eventual Selection of a High Fitness Proliferative Cell State

To understand the functional consequences of the observed transcriptional changes beyond the changes in net growth rates and cell spreading that have been previously reported experimentally (13), we quantified pathway-level activity by performing single-sample Gene Set Enrichment Analysis (ssGSEA) on the PID (Pathway Interaction Database) pathway set. A Principal Component Analysis (PCA) based on these pathway scores (**Fig. 2A**) recapitulated the strong separation seen at the gene level. PC1, explaining 53.57% of the variance, corresponded to the ’Ancestral-Selected’ axis, while PC2, explaining 24.12% of the variance, represented the ’Stiff-Soft’ axis. This confirms that the evolutionary selection process, rather than the initial stiffness cue, drives the most substantial functional cell state enrichment. We therefore sought to compare the pathway-level changes across two distinct selection pressures that enrich for differently selected cell variants (1) selection acting on cells maintained on a stiff substrate (Stiff Ancestral to Stiff Selected) and (2) selection within a soft environment, which involves an initial stiffness change (proxied by a comparison between Stiff Ancestral and Soft Ancestral conditions) followed by long-term selection (Soft Ancestral to Soft Selected).

**Figure 2.**
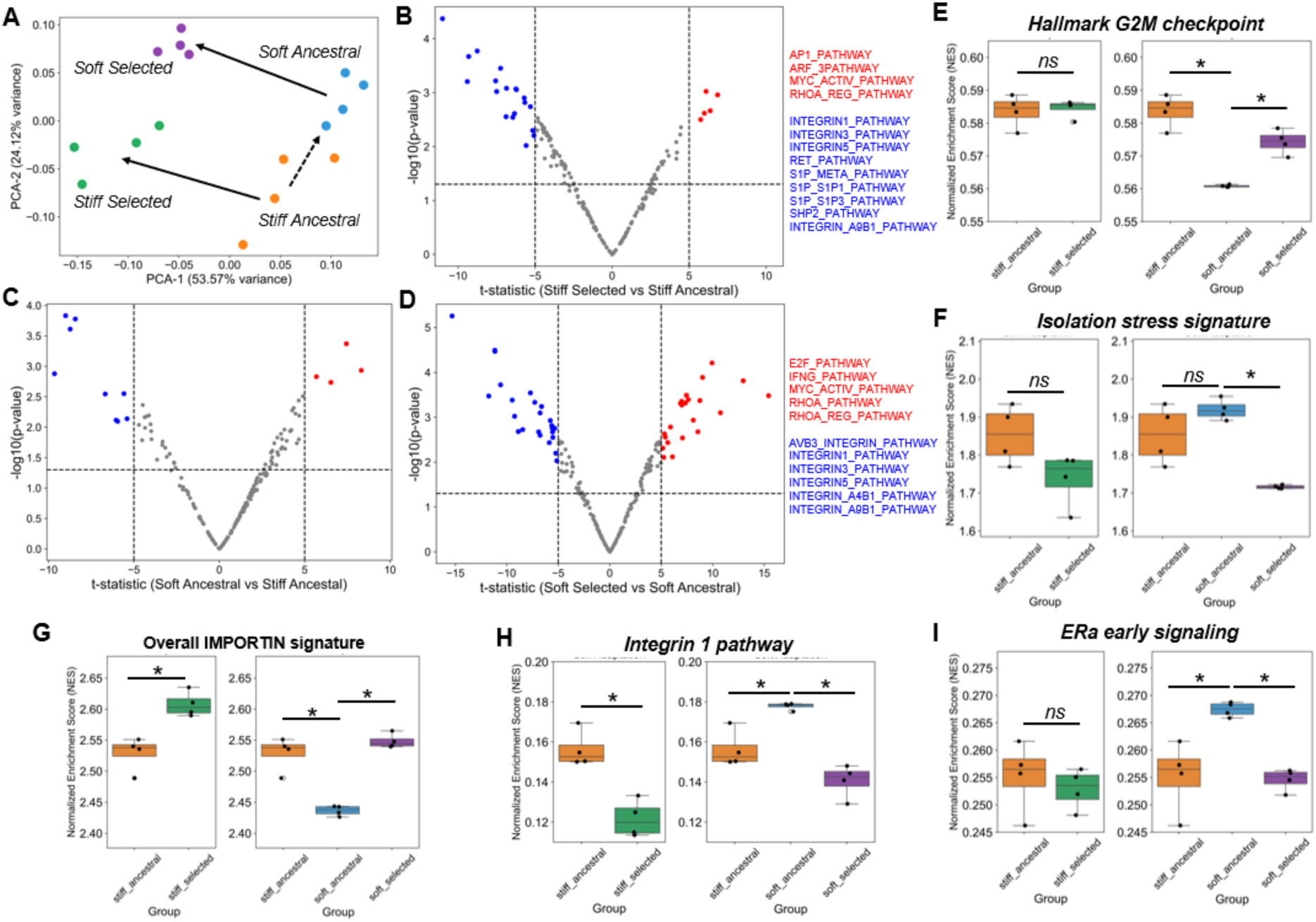
Pathway-Level Analysis Reveals Acute Mechanosensitive Stress and Distinct Phenotypic Variations. **(A)** Principal Component Analysis (PCA) of PID pathway ssGSEA scores for all RNA-seq samples. PC1 (53.57% variance) delineates the Ancestral-Selected axis, while PC2 (24.12% variance) distinguishes the Soft-Stiff axis. **(B)** Volcano plot showing differentially active PID pathways between Stiff Selected and Stiff Ancestral cells. **(C)** Volcano plot illustrating differentially active PID pathways between Soft Ancestral and Stiff Ancestral cells. **(D)** Volcano plot showing differentially active PID pathways between Soft Selected and Soft Ancestral cells. **(E)** ssGSEA scores for the HALLMARK_G2M_CHECKPOINT pathway across representative conditions. **(F)** ssGSEA scores for an “Isolation Stress” pathway signature. **(G)** ssGSEA scores for overall Importin signature. **(H)** ssGSEA scores for Integrin1 signaling pathway. **(I)** ssGSEA scores for ERα signaling pathway. Asterisks indicate statistical significance based on Students t-test with p-value < 0.05.

We first examined the stiff-substrate specific phenotypic selection process. A volcano plot comparing pathway scores between Stiff Selected and Stiff Ancestral cells (**Fig. 2B**) revealed a significant upregulation of pathways including c-Myc transcriptional activation and regulation of RhoA activity. Concurrently, many integrin-related pathways were significantly downregulated. This trend suggests that prolonged selection in a consistently stiff environment favors cells with heightened oncogenic signaling and cytoskeletal regulation while reducing reliance on the adhesion-mediated signaling pathways. This observation is consistent with our previous experimental work demonstrating Rho-regulated cell spreading as a selected trait (13).

We next investigated the soft-substrate selection route. The initial transfer on to the soft substrate (i.e. from Stiff Ancestral to Soft Ancestral) (**Fig. 2C**) induced fewer significant pathway alterations, consistent with our PCA findings that the selection axis (PC1) accounts for the majority of transcriptomic variance. Among the pathways that were altered, c-Myc transcriptional activation, aurora kinase A/B pathways were significantly downregulated in Soft Ancestral cases, indicating a drop in cell proliferation consistent with a lower reported growth rate of the soft ancestral cells compared to stiff ancestral cells (13). However, the subsequent selection of the Soft Selected cell state from the Soft Ancestral cell population (**Fig. 2D**) was associated with more pronounced differences in pathway activities. Similar to the stiff selected cells, the RhoA activity pathway was significantly upregulated. Crucially, the E2F transcription factor pathway was also upregulated considerably, providing a direct molecular link to the enhanced cell proliferation observed experimentally (13,15). The interferon gamma (IFN-γ) signaling pathway was also increased, and similar to the stiff-selected state, multiple integrin pathways were downregulated upon selection (**Fig. 2D**).

To further corroborate the link between selection on soft substrate and the enrichment of a high proliferative cell state, we specifically tracked the Hallmark G2M checkpoint pathway from MSigDB (**Fig. 2E**). This pathway activity score was uniquely and significantly downregulated in the Soft Ancestral state when compared to the Stiff Ancestral population, aligning with the experimental observations of reduced proliferation upon stiffness reduction (13). This activity was fully restored in the Soft Selected state to levels comparable to stiff-cultured cells, strongly supporting the hypothesis that the initial encounter with a soft substrate enriches for a “stressed” and “low-fitness” state marked by cell cycle attenuation, which is subsequently resolved when higher fitness cell clones are selected for. Similar trends were observed for aurora kinases (AURKA, AURKB) gene expression profiles, which are also known to be crucial for cell division (**Supplementary Fig. S1A-B**). Experimentally, it has been demonstrated that the soft ancestral cells are unable to spread on the soft substrate, and feature rounded, irregular Lamin B1-stained nuclei, show an absence of central filamentous actin (F-actin) stress fibers, and reduced nuclear YAP (13). We investigated whether this phenotype is a form of “isolation stress” as has been previously reported (16–18). Indeed, an “isolation stress” signature comprised of FN-binding integrins αvβ3 and α5β1 was found to be highly active in soft ancestral cells and was significantly downregulated in the soft selected state (**Fig. 2F**). Furthermore, the hypoxia pathway (MSigDB Hallmark Hypoxia) was found to be upregulated explicitly in soft ancestral cells (**Supplementary Fig. S1C**), which is known to be high in such type of isolation stress (18) and in the scenarios of anoikis resistance (19).

This acute stress/low fitness state is linked to several compensatory responses for cell survival. Of note, we observed a global drop in the importin signature, encompassing all importins, in the Soft Ancestral state (**Fig. 2G**). More specifically, the expression of importin 7 (IPO7), the major YAP1-specific importin (20), was downregulated in the soft ancestral cells (**Supplementary Fig. S1D**). This trend is in line with YAP1 retention in the cytoplasm as observed experimentally (13). Soft ancestral cells, which appear rounded and are unable to spread, indicative of poor matrix engagement, exhibit a brief upregulation of integrin signaling pathways in response to reduced substrate stiffness (**Fig. 2H**). This response does not persist, as long-term selection ultimately enriches cells that can cope with soft conditions without relying on this temporary integrin activation (**Fig. 2D, 2H**). Furthermore, we observed a significant upregulation of ERα signaling (**Fig. 2I**). While these are triple-negative breast cancer (TNBC) cells, an upregulated ERα signaling may point to a broader, stress-induced upregulation of nuclear hormone receptor signaling similar to prior observations in prostate cancer cells with associated epigenetic changes when grown on different stiffness conditions (21). This acute stress response was also marked by a significant metabolic shift. Expression levels of RXRA as well as the RXR-VDR pathway were found to be highest in the Soft Ancestral state (**Supplementary Fig. S1E-F**). These metabolic programs are subsequently downregulated following selection, which specifies a broad metabolic signature of the ’brown’ ancestral module (**Fig. 1F**) as a short-term survival response in the fitness-compromised state on soft ECM.

### Selection on Soft Substrate Involves a Reduction in Chromatin Accessibility of the Ancestral Metabolic Program

To investigate whether the observed transcriptional differences between the abovementioned four experimental groups are explained by differences in their corresponding chromatin landscapes, we performed ATAC-seq (Assay for Transposase-Accessible Chromatin using sequencing) on them. We focused our analysis on the genes identified in the WGCNA modules and quantified the overlap between differentially expressed genes (DEGs) and genes with differentially accessible (DA) chromatin peaks within their promoter regions (±1 kbp from the gene’s start site). We compared three principal stages: the ancestral stressed, low-fitness state on soft ECM (Soft Ancestral vs Stiff Ancestral, **Fig. 3A**), the subsequent soft-substrate selected cell states (Soft Selected vs Soft Ancestral, **Fig. 3B**), and the corresponding stiff-substrate selected cell states (Stiff Selected vs Stiff Ancestral, **Fig. 3C**). Consistent with our transcriptomic data, the initial soft substrate specific low fitness cells (**Fig. 3A**) resulted in substantially fewer genes with correlated differential expression and accessibility compared to long-term selection. Consequently, cells that persist through selection exhibit far stronger accessibility relative to Soft Ancestral than those observed between Soft Ancestral and Stiff Ancestral. In both the Soft selected and Stiff selected conditions (**Fig. 3B, 3C**), the upregulated blue and black module genes showed a high incidence of associated promoter accessibility changes compared to respective ancestral groups. However, the direction of this change was not uniform; for the upregulated genes in these modules, approximately 55% of them had a higher promoter accessibility, while 45% surprisingly had a decrease. This complex relationship suggests that while chromatin accessibility is involved, it is not the sole contributor to these programs in the selected cells. This pattern is consistent with previous work demonstrating changes in chromatin accessibility need not be always concordant with transcriptional changes (22). In contrast, the downregulated genes in the brown “ancestral” module in selected cells compared to respective ancestral groups revealed a more direct epigenetic link. Across both the selection scenarios, a substantial majority (∼60-65%) of the downregulated brown module genes also exhibited a significantly lower promoter accessibility (**Fig. 3B, 3C**). This strong concordance between RNA-seq and ATAC-seq measurements indicates that epigenetic silencing of the ancestral gene program is a defining feature of the selected state. This trend is most pronounced in the Soft Ancestral versus Soft Selected comparison and is further supported by ATAC-seq average promoter accessibility profiles and heatmaps, which show a clear skew toward reduced chromatin accessibility at promoters (±1 kb from the TSS) of downregulated brown module genes (**Fig. 3D**).

**Figure 3.**
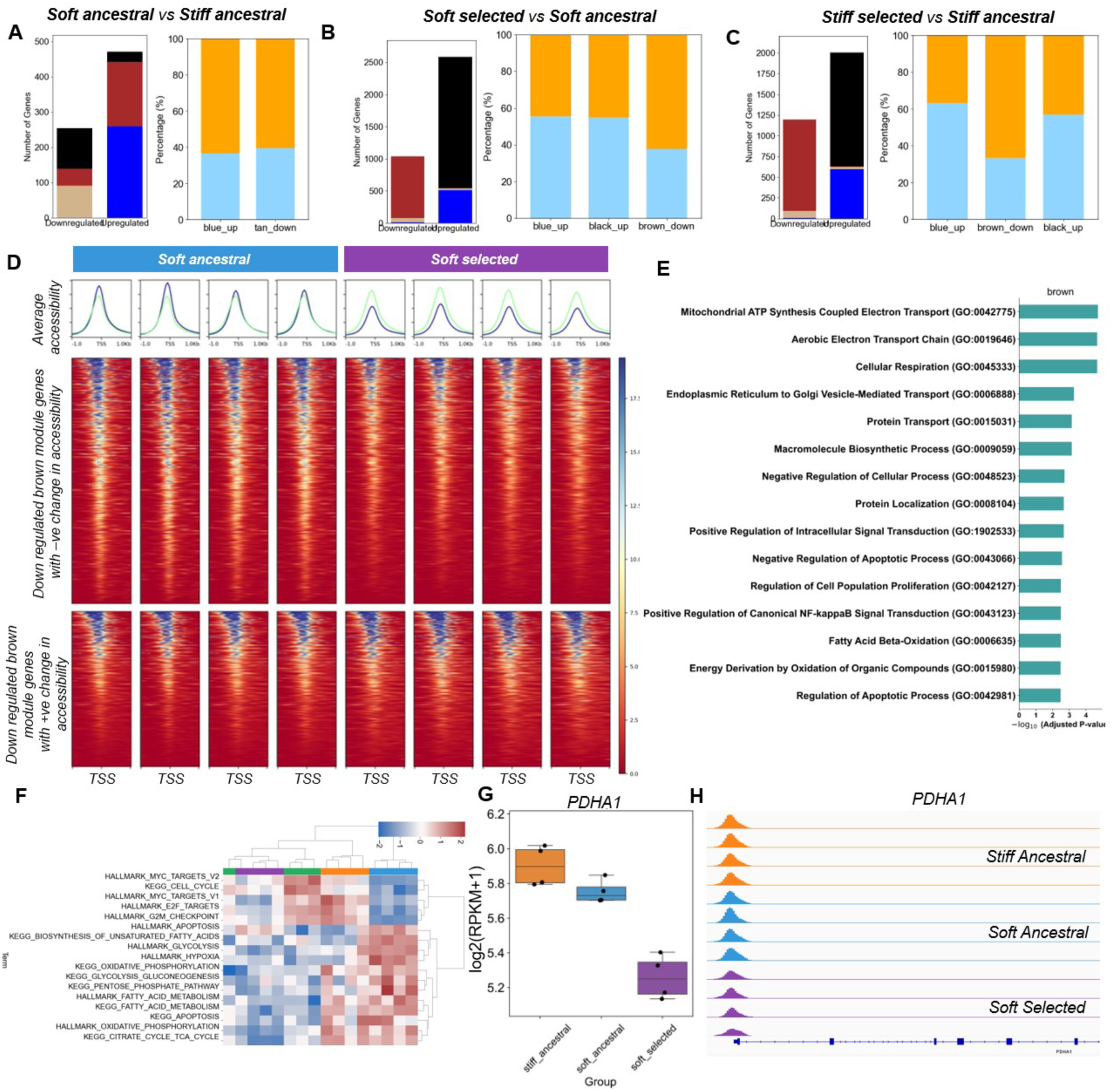
Phenotypic Difference During selection is Associated with Epigenetic Silencing of the Ancestral Program via Reduced Chromatin Accessibility. (A-C) Stacked bar plots on the left depict the number of module genes (blue, black, brown, tan) that are upregulated or downregulated along with corresponding changes in promoter accessibility (±1 kbp from the gene start). The bar plots on the right show the number of differentially expressed module genes exhibiting increased (sky blue) or decreased (orange) promoter accessibility. Each comparison is shown for **(A)** Stiff Ancestral vs. Soft Ancestral, **(B)** Stiff Ancestral vs. Stiff Selected, and **(C)** Soft Ancestral vs. Soft Selected. Percentages highlight the strong skew toward decreased accessibility for downregulated brown module genes in selection trajectories (B, C). (**D)** Representative ATAC-seq average promoter (±1 kb from transcription start site, TSS) accessibility profiles and corresponding heatmaps for downregulated brown module genes in soft ancestral and soft selected states. Average accessibility profiles are shown for genes exhibiting negative (blue) and positive (green) changes in accessibility across individual replicates. Heatmaps depict promoter accessibility of the corresponding gene sets, ordered by accessibility signal. A predominant concordant trend is observed, with a larger fraction of downregulated brown module genes displaying reduced promoter accessibility. **(E)** Gene ontology (GO) analysis for the subset of brown module genes that are both transcriptionally downregulated and show decreased promoter accessibility. Enriched terms include metabolic pathways. **(F)** Clustered heatmap showing z-score–normalized ssGSEA enrichment scores for Hallmark and KEGG pathways related to metabolic and cell cycle processes across the indicated conditions: Stiff Ancestral (orange), Soft Ancestral (blue), Soft Selected (purple), and Stiff Selected (green) **(G)** Gene expression levels for PDHA1 gene across the soft selection trajectory compared to stiff ancestral cells. **(H)** Representative ATAC-seq genome browser tracks for PDHA1 gene. The three conditions in order are Stiff Ancestral, Soft Ancestral, and Soft Selected cells.

We next performed Gene ontology (GO) analysis for genes exhibiting reduced expression as well as promoter accessibility in Soft selected vs. Soft ancestral cells and observed diverse metabolism-related pathways (**Fig. 3E**). Further pathway analysis using ssGSEA scores for KEGG and MSigDB Hallmark metabolic pathways revealed that soft ancestral cells were more enriched for oxidative phosphorylation, fatty acid biosynthesis and metabolism, and the pentose phosphate pathway. On the other hand, they expressed lower levels of cell cycle and associated pathways (**Fig. 3F**). Overall, we observed a significant downregulation of the brown module gene PDHA1 (Pyruvate dehydrogenase E1 subunit alpha 1) (**Fig. 3G**). This gene is a crucial component of the pyruvate dehydrogenase (PDH) complex, which facilitates oxidative phosphorylation (OXPHOS) by converting pyruvate into acetyl-CoA. This conversion allows acetyl-CoA to enter the tricarboxylic acid (TCA) cycle (23). This alteration was also accompanied by a marked reduction in promoter accessibility of PDHA1 gene in soft-selected cells compared to both stiff ancestral and soft ancestral cells. (**Fig. 3H**). Together, these findings reveal that phenotypic selection involves coordinated chromatin-level repression of the ancestral brown module metabolic program, as characterized by reduced promoter accessibility and expression of oxidative metabolism genes such as PDHA1.

### Selection on the Soft Substrate Involves De Novo DNA Methylation to Stably Silence the Ancestral Program

To assess an additional layer of regulation beyond chromatin accessibility in the Soft Selected cells, we performed DNA methylation profiling using Reduced Representation Bisulfite Sequencing (RRBS). Soft Ancestral cells, compared to Stiff Ancestral ones, exhibited far fewer DMCs compared to Stiff Ancestral cells, with only ∼2.5% of CpG sites significantly altered (1.26% hypermethylated and 1.32% hypomethylated) (**Fig. 4A**). Strikingly, the process of selection amplifies the differences in DMCs – we noticed much higher DMCs in comparing the scenarios of a) Stiff Ancestral vs. Soft Ancestral with either b) Stiff Selected vs. Stiff Ancestral (∼9% significantly dysregulated CpG methylations) or with c) Soft Selected vs. Soft Ancestral (∼6.76% significantly dysregulated CpG methylations) (compare **Fig 4A** with **Fig 4B – i, ii**). Notably, in both the selection scenarios, we observed a significantly greater number of hypermethylated DMCs than hypomethylated ones (soft selection: 4.98% hypermethylated and 1.77% hypomethylated; stiff selection: 6.73% hypermethylated and 2.28% hypomethylated). This skew towards a net gain of methylation strongly suggests the involvement of a widespread change in *de novo* methylation in both stiff and soft selected cells, potentially contributing to their higher fitness.

**Figure 4.**
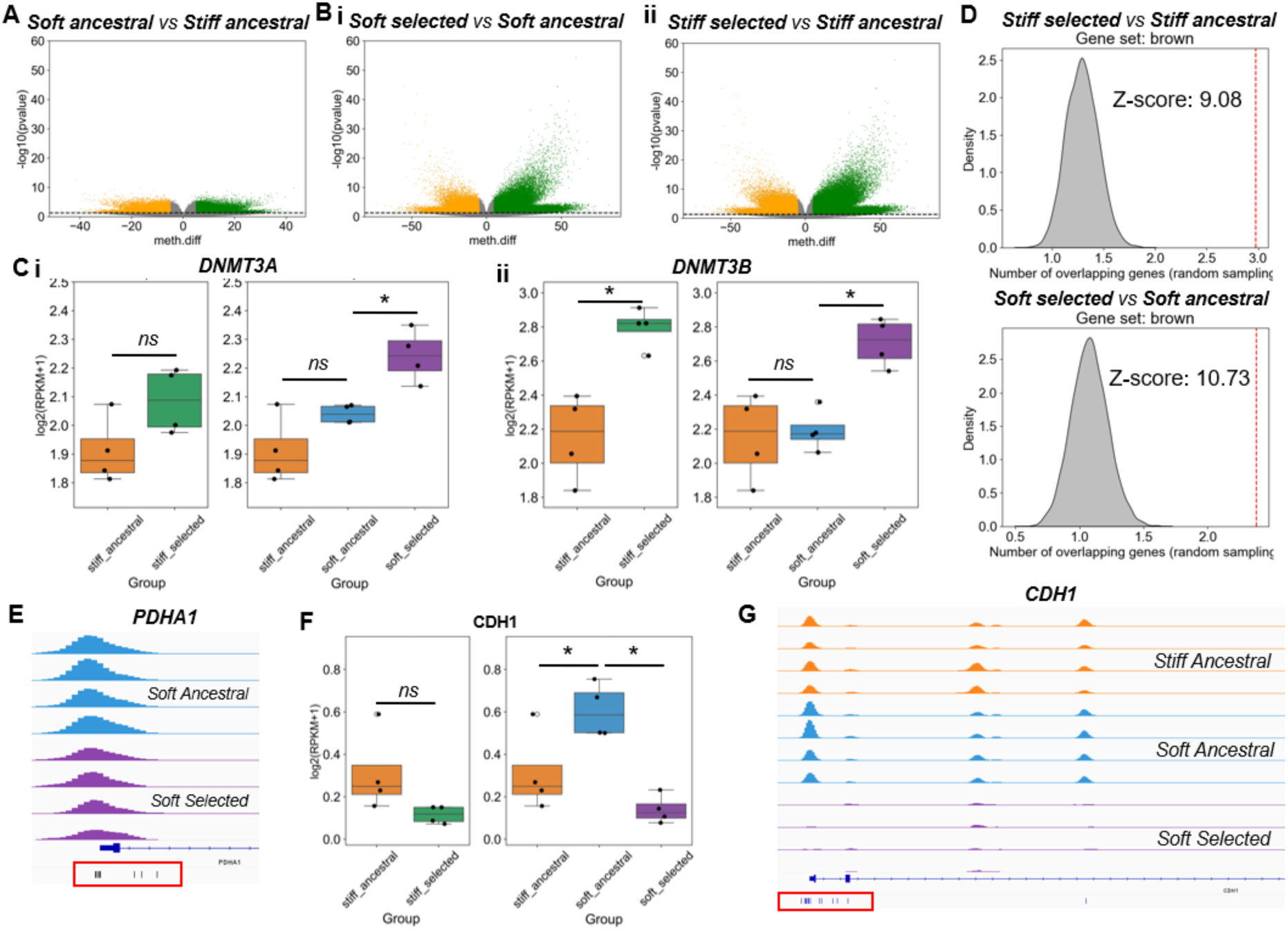
Analysis of DNA Methylation Changes Associated with Soft Selection. **(A)** Volcano plots of DMCs for Soft Ancestral vs. Stiff Ancestral cells. (p < 0.05 and |methylation difference| > 0) **(B)** Volcano plot of Differentially Methylated CpGs (DMCs) comparing (i) Soft Selected vs. Soft Ancestral and (ii) Stiff Selected vs. Stiff Ancestral cells. Green: hypermethylated CpGs; orange: hypomethylated CpGs. (p < 0.05 and |methylation difference| > 0) **(C)** Normalized gene expression of (i) DNMT3A and (ii) DNMT3B for stiff selection and soft selection. Asterisks indicate statistical significance. (**D)** Enrichment analysis of brown module genes with promoter hypermethylation clusters (>10 DMCs, <100 bp spacing, ±1 kbp from gene start). Histograms show the empirical null distribution (chromosome-controlled random sampling). The vertical line indicates the observed frequency for brown module genes, with Z-scores shown for Stiff Selected (vs. Stiff Ancestral) and Soft Selected (vs. Soft Ancestral) comparisons. **(E)** Genome browser tracks showing ATAC-Seq accessibility and RRBS methylation profiles for PDHA1 gene for Soft Ancestral and Soft Selected cells. **(F)** Bar plot of normalized CDH1 expression across conditions. **(G)** Genome browser tracks showing ATAC-seq (accessibility) and RRBS (methylation) data at the CDH1 promoter for four replicates each of stiff ancestral (orange), soft ancestral (blue) and soft selected (purple).

To determine the molecular machinery contributing to the methylation patterns observed in both the selected populations, we examined the expression of DNA methyltransferases. Consistent with a surge in *de novo* methylation, the expression levels of DNMT3B – an enzyme responsible for establishing new methylation patterns (24) – was significantly upregulated in both the selected samples compared to their ancestral counterparts (**Fig. 4C, i-ii**). In contrast, the expression of DNMT1, the maintenance methyltransferase that copies existing methylation patterns during DNA replication (25), followed a different pattern. When compared with corresponding ancestral populations, it was down regulated in the Soft Ancestral state, concordant with the transient cell cycle arrest and reduced proliferation, but was higher in the Soft Selected state, paralleling the increased cell proliferation and higher localization of YAP1 to the nucleus (**Supplementary Fig. S2A**).

We next sought to identify the specific gene programs targeted by this extensive hypermethylation. We queried the promoter regions (±1 kbp from the gene start) of genes belonging to all four WGCNA modules for the presence of hyper-methylated CpG clusters rather than individual CpGs. Such clusters – a hallmark of stable gene silencing – are defined as having at least 10 hypermethylated CpGs with no more than 100 base pairs separating them. To gain more functional relevance, we included only those genes that were differentially expressed and contained at least one such hypermethylated cluster. We then compared the observed frequency of these events in each module against an empirical null distribution, generated by chromosome-controlled random sampling of genes, to determine statistical over-representation. This analysis yielded a key result: the brown “ancestral” module genes were highly over-represented among genes acquiring promoter hypermethylation in both Soft Selected and Stiff Selected cells compared to their ancestral counterparts (Z-score > 9 for both soft and stiff selected) (**Fig. 4D**). This finding offers direct evidence that a subset of the ancestral program, characterized by high metabolic activity, is not merely transcriptionally downregulated, but also is actively silenced through targeted *de novo* DNA methylation of its promoters for potential longer timescales. For example, PDHA1, which was differentially accessible (**Fig. 3G**) as well as significantly downregulated (**Fig. 3H**) in soft selected cells, was also hypermethylated in soft selected cells compared to soft ancestral cells (**Fig. 4E**).

Furthermore, we identified critical genes within the brown module that exemplified the *de novo* silencing mechanism and how it might affect the cell cycle status of cells. CDH1 (E-cadherin), a key epithelial marker, also known to be critical in the progression of cell cycle (26), was found to be highly expressed in Soft Ancestral state as compared to both Soft Selected and Stiff Ancestral state (**Fig. 4F**). This transcriptional repression was correlated with its promoter both becoming inaccessible (**Fig. 4G**) and acquiring dense hypermethylation (**Fig. 4G**, red box). This combination of chromatin closure and DNA methylation ensures a robust and potentially heritable silencing of its expression. Similarly, the YAP1-mediated pro-apoptotic gene TAp73 (27) was markedly upregulated during the Soft Ancestral state, which was lower in the Soft Selected cells, concordant with changes in terms of hypermethylated CpG clusters at the TAp73 promoter (**Supplementary Fig. S2B-C**).

Collectively, these results demonstrate that epigenetic silencing is a characteristic feature of soft-selected cells. Thus, under long-term selection, genetic clones that silence stress-response and pro-death genes by co-opting the *de novo* methylation machinery can execute a heritable, pro-survival, and highly proliferative phenotype.

### A Coupled Mechanotransduction and Cell Cycle Gene regulatory Network to Model Changes in YAP Localization and Proliferation Changes on Soft Substrates

To understand the dynamic coordination between mechanosensing, change in cell growth rate, and the fitness advantage due to epigenetic marks on selected genetic clones, we constructed a mechanistic mathematical model integrating our observations with previously reported experimental inter-connections among the key molecular determinants identified so far. This model incorporates these distinct biological processes to explain how genetic variation, along with epigenetic regulations, can drive a coordinated pattern of gene expression, enabling high-fitness, proliferative cell states on soft substrates.

MDA-MB-231 cells growing on ultra-stiff substrate (glass/plastic), as closely mimicked by Stiff Ancestral cells, are defined by a basal program that involves YAP1 localized to the nucleus with extensive cell spreading and high cell proliferation (13). Prior work on cancer cells on two-dimensional (2D) surfaces has indicated that nuclear flattening in spread cells promotes YAP/TAZ nuclear localization (28). Furthermore, lowering substrate stiffness has been shown to lead to the loss of nuclear YAP and result in reduced YAP-dependent cellular proliferation (11,29). Thus, it remains unclear how certain genetic clones can demonstrate cell spreading, exhibit subsequent YAP translocation to the nucleus, and high proliferation. Through a mechanism-based mathematical model, we hypothesize that a small number of genetic clones might possess molecular mechanisms to allow spreading and YAP translocation to the nucleus, thus creating a proliferation-permissive state. Furthermore, epigenetic alterations, such as CDH1 promoter hypermethylation in these clones, can further reduce cell cycle duration times and confer greater fitness.

Our proposed gene regulatory network model is composed of two key parts: i) a mechanotransduction module that models how YAP nuclear localization is dependent on external stiffness, and the interplay of this module with FAK and RhoA signaling, and ii) a cell cycle module that models how the cell cycle duration depends on key regulators such as CDH1 and how epigenetic silencing of high levels of CDH1 can reduce cell cycle duration. The mechanotransduction module for our ordinary differential equation (ODE) system is adapted from a previously established model by Scott *et al.* (29), which provides a detailed quantitative description of how ECM stiffness is sensed at focal adhesions and translated through FAK and RhoA signaling. It ultimately controls the nuclear localization of the YAP/TAZ co-activator as a function of nuclear pore stretching and Lamin A activation. In this study, we demonstrated that the levels of most importin molecules, and more specifically the expression of IPO7 (**Fig. 2G, Supplementary Fig. S1D**), the primary carrier for YAP1 (20), differ significantly across the various experimental conditions. We also considered the feedback activation of RhoA levels by YAP1 through RacGAP1 – a potential activator of RhoA (30) that was significantly low in Soft Ancestral cells as compared to both Soft Selected and Stiff Ancestral cells (**Supplementary Fig. S3A**).

For the cell cycle module, we adapted a mathematical model by Charan *et al*. (31) that models cell cycle oscillations and duration through a gene regulatory network composed of CDH1, CDC20, and CycB (CCNB1). Here, based on prior literature (11,32,33), we assume that CDC20-based activation of the cell cycle and completion is MYBL2-dependent, which itself is regulated by YAP1 (11). This YAP/MYBL2/CDC20 cascade establishes a direct, stiffness-dependent link that controls the cell proliferation rate. Thus, in our framework, both the YAP localization to the nucleus, YAP activation of MYBL2 and the optimal levels of cell cycle regulators would be crucial to drive the cell cycle successfully. CDH1 upregulation has been shown to lead to cell cycle arrest in breast cancer cells (34). In this study, we have observed that CDH1 expression is high in Soft Ancestral cells, which can potentially explain the lower growth rate (i.e. cell cycle of longer duration) in these cells (**Fig. 4F**).

Experimentally, the Soft Selected state involves the assembly of central filamentous actin (F-actin) stress fibers, which were absent in soft ancestral cells (13). We observed upregulation of FAK (PTK2) –a molecule upstream of RhoA/YAP signaling cascade (35) – levels in the soft ancestral and soft selected cells when compared to the stiff ancestral population (**Supplementary Fig. S3B**). To represent the mechanosignaling components upon a change of substrate as observed in specific genetic variants, we consider a variable that accounts for upregulation of total FAK levels and subsequent localization of YAP1 to the nucleus, as a function of F-actin levels (35,36). Given that FAK inhibitors can cause enhanced proteasomal degradation of DNMT3 molecules (37), this variable can also indirectly activate genes such as DNMT3. Finally, to model the epigenetic silencing of the ancestral program, as observed in our multi-omics analysis, specifically for the CDH1 gene, we introduced a variable that accounts for the level of epigenetic memory to model the effects of upregulated DNMT3 empirically. This variable, thus, acts as a proxy for *de novo* DNA methylation and can lead to transcriptional suppression of CDH1 in the fitter genetic variants. In principle, both variables are heritable and are potentially encoded by the underlying genetic differences. To account for genetic heterogeneity in our virtual cell population, most values of the model parameters are sampled from a Gaussian distribution (see **Supplementary Methods**). This integrated gene regulatory network provides a comprehensive molecular description to explain how certain genetic clones can exhibit higher fitness on soft substrates through gene expression and epigenetic changes.

### A Mechanistic Population Model Recapitulates the Selection of High-Fitness Genetic Clones on Soft Substrates Driven by Gene Expression and Epigenetic Differences

To explain the observed molecular patterns during the soft selection process in a quantitative, coherent framework, we propose a regulatory network that captures the dynamic interplay between mechano-transduction, cell cycle control, and epigenetic memory in genetically diverse cell populations (**See Supplementary Methods**). First, we establish the baseline behavior of this model, which is built on prior formulations (29,31). This model recapitulates fundamental mechano-sensitivity of the YAP/TAZ pathway, showing a sigmoidal relationship where the steady-state nuclear-to-cytoplasmic ratio of YAP is high on stiff substrates and low on soft substrates (**Fig. 5A**).

**Figure 5.**
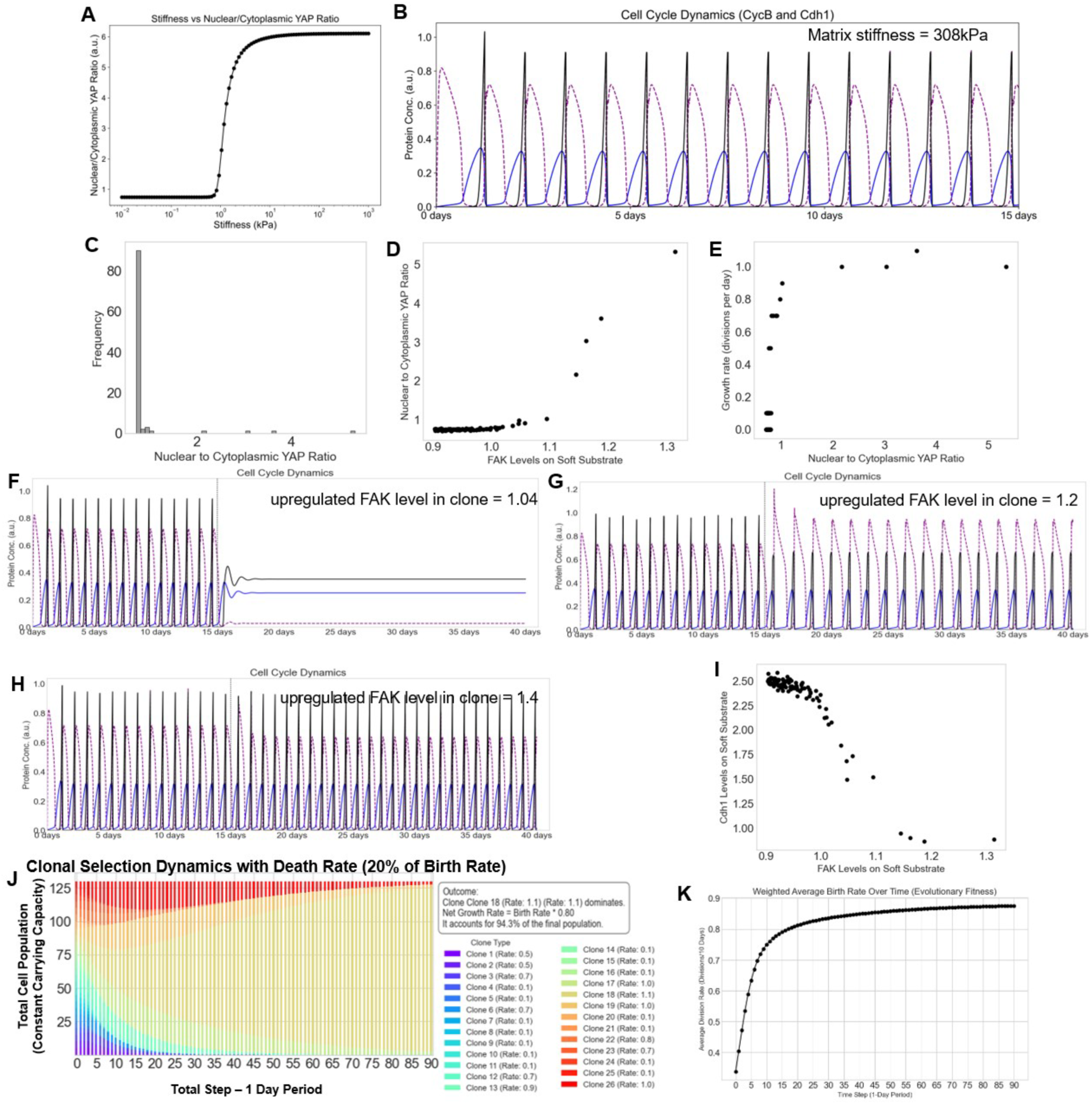
A Coupled Mechanotransduction and Cell Cycle Model Recapitulates Selection of Fit Genetic Clones. **(A)** Model steady-state analysis showing the simulated steady-state nuclear-to-cytoplasmic ratio of YAP as a function of increasing ECM stiffness, demonstrating the model’s core sigmoidal mechanosensitivity. **(B)** Simulation on high stiffness (308 kPa) shows robust, periodic oscillations of CycB (blue), CDH1 (purple), and CDC20 (black), corresponding to a stable cell cycle duration of ∼25 hours. **(C)** Distribution of nuclear to cytoplasmic YAP ratio of Soft Ancestral cells. Note rare clones have a large nuclear to cytoplasmic ratio. **(D)** Scatterplot showing the upregulated FAK levels in a subset of cells which also express higher nuclear to cytoplasmic YAP ratio. **(E)** Scatterplot showing the nuclear to cytoplasmic YAP ratio and the growth rate of individual cells (calculated from cell cycle duration) showing a positive correlation. Representative simulations of different cancer cell clones on high stiffness (308 kPa) followed by low stiffness (3 kPa) shows robust, periodic oscillations of CycB (blue), Cdh1 (purple), and CDC20 (black), in stiff substrate but **(F)** cell cycle arrest for clone not upregulating FAK substantially, **(G)** longer cell cycle duration of a clone with intermediate FAK upregulation on soft substate and **(H)** 23 hour fast cell cycle duration with highest levels of FAK upregulation. **(I)** Scatterplot showing the relationship between upregulated FAK levels in different cancer cell clones due to different extents of epigenetic silencing. **(J)** In-silico selection experiment showing the gradual takeover of the fittest clone (clone 18) over a 90-day selection period. **(K)** Population weighted growth rate average of cancer cells over the 90-day selection period.

Next, we consider that initial cells grow at a stiffness of 308kPa, leading to a nuclear to cytoplasmic ratio ∼4.5, which leads to a constant cycling of cells, as is demonstrated by oscillations in the levels of CycB (CCNB1), CDH1 and CDC20 with a cell cycle duration of approximately 25 hours (**Fig. 5B**). Next, to mimic the experimental setup of substrate change, we simulated a sudden stiffness change from 308kPa to 1kPa at the 15-day mark. This caused a sudden drop in the nuclear to cytoplasmic YAP ratio to a value <1 in a majority of cells, with a small subset of parameter sets showing higher nuclear to cytoplasmic YAP ratio >2 (**Fig. 5C**). This increase in nuclear to cytoplasmic YAP levels in a small subset of cells was due to the upregulation of FAK in these cells (**Fig. 5D**). Consequently, these cells had a higher fitness on soft substrates as demonstrated by a faster growth rate (**Fig. 5E**). Cells which could not upregulate FAK and localize YAP to the nucleus generally exhibited a state of cell cycle arrest (**Fig. 5F**). Intermediate levels of FAK upregulation led to a continuation of cell cycle but with a longer duration of cell cycle of nearly 36 hours (growth rate ∼0.67 day-1) (**Fig. 5G**). However, cell clones which upregulated FAK the highest maintained a cell cycle duration of nearly 23 hours (growth rate ∼1.03 day-1) (**Fig. 5H**). The highest levels of FAK upregulation were also associated with lower levels of CDH1 (**Fig. 5I**), highlighting the role of epigenetic silencing of CDH1 to modulate the cell cycle duration. Finally, we performed an *in-silico* selection experiment on different genetic clones that had a non-zero growth rate (only cells with a non-zero growth rate could establish clonal colonies). We observed that the most dominant clone, i.e., the clone which upregulated FAK and downregulated CDH1 to the most significant extent, was the cell which eventually took over the population (clone 18 with a growth rate of 1.1) (**Fig. 5J**). Further, the average cell growth rate, weighted by their abundance, showed a monotonic increase from 0.4 day-1 to 0.85 day-1 within 20-25 days of the start of the selection experiment, recapitulating our experimental observations (13) (**Fig. 5K**).

## Discussion

Despite extensive knowledge about the YAP-mediated transcriptional responses to ECM stiffness (10,38), the contribution of mechanotransduction pathways to long-term selection in soft ECM environments remains unclear. Soft microenvironments, often present at metastatic sites, can induce chemoresistance (39) and stem-like behavior (40). Thus, investigating the molecular and functional traits associated with long-term selection on soft ECM environments can hold the key for potential anti-metastatic strategies. We address this gap by providing a multi-omics mechanistic dissection of the evolution of breast cancer cells under prolonged culture on soft ECM. This integrative approach allows us to delineate molecular determinants that shape long-term selection in soft mechanical environments.

Prior studies have shown that the concordance between chromatin accessibility and gene expression profiles can be low (22). Transcriptional bursting and noise in gene expression can often decouple these layers of gene expression regulation. Open chromatin often precedes transcription in a poised state (41), and stochastic transcriptional bursting (42) can generate substantial heterogeneity in mRNA levels independent of a stable chromatin architecture. Our observations of molecular determinants of Soft Selected cells support the fact that chromatin accessibility is not strictly coupled with gene expression overall. However, a tight coordination observed here between promoter hypermethylation, reduced accessibility and transcriptional silencing for specific gene products such as CDH1 suggests that selection of specific genetic clones favors high-fitness variants with robust, hard-wired genomic state potentially minimizing the impact of stochastic noise that can drive spontaneous cell-state transitions (43–45).

Evolutionary selection eventually enriches specific variants that possess higher proliferative fitness. Through multi-omics analysis, we hypothesize that these genetic clones can upregulate FAK that facilitates YAP nuclear localization which is also concordant with cell spreading (13,46). These selected variants activate DNMT expression to suppress potential obstacles to cell cycle progression, such as high levels of CDH1, and demonstrate strong activation of the YAP-MYBL2 axis, which is also known to be the central regulator of mitotic gene expression acting through Myb–MuvB complexes (11). Beyond increased cell cycle activity, these cells exhibit a coordinated repression of the metabolic and stress-response programs (brown module) that dominated the ancestral state, a finding supported by ATAC-seq data revealing reduced promoter accessibility within the brown module of the soft-selected population.

Literature on metastatic organotropism suggests that successful colonization of secondary metastatic sites with distinct rigidity – such as the mechanically softer brain tissue versus the rigid niche of the bone – is contingent upon cells possessing specific metabolic phenotypes (47). Our findings imply that these distinct metabolic requirements are not solely defined by nutrient availability at metastatic sites but also may be enforced by tissue mechanics. Consequently, the Soft Selected phenotype represents a subpopulation whose metabolic state was potentially compatible with growth on a soft matrix, allowing them to overcome metabolic bottlenecks that may restrict metastatic outgrowth in softer organs such as the brain (47,48).

Our computational work opens several critical avenues for future investigation, particularly regarding the direct validation of the model’s predictions. For instance, treatment of “Soft Ancestral” cells with DNMT inhibitors is predicted to prevent the epigenetic silencing of CDH1; while this should still allow cells to proliferate, it is likely to lock them in a slow-cycling phase. Conversely, the application of FAK inhibitors could potentially abrogate the selection process on soft substrates by causing cell cycle arrest at the G2/M phase. Additionally, a MYBL2 knockdown is expected to selectively arrest the “Soft Selected” cells, confirming the essential role of this regulator in the selected state. Our findings indicate the critical role of FAK expression in enabling higher cell proliferation for the Soft Selected cells, thus administering FAK inhibitors (49) can be a potent therapeutic strategy to prevent the selection of specific genetic clones with higher fitness in soft ECM environments.

## Materials & Methods

### RNA-Seq data analysis

RNA-seq data for MDA-MB-231 ancestral and stiffness-selected cells cultured on soft (1 kPa) and stiff (308 kPa) substrates were obtained from GSE255829. FPKM values were downloaded from NCBI GEO and log₂(FPKM + 1) normalization was applied. Differential gene expression between sample groups was determined using a Student’s t-test with a significance threshold of *p* < 0.05.

### Principal Component Analysis

Principal Component Analysis (PCA) was performed on normalized gene expression values using the scikit-learn Python package to identify major sources of variance across samples. The expression matrix was mean-centered and scaled prior to PCA, and the top principal components were used to visualize sample clustering and assess variability between conditions.

### WGCNA analysis

Weighted Gene Correlation Network Analysis (WGCNA) was performed to identify gene co-expression modules. log₂(FPKM + 1) normalised counts were used as input. Low-quality genes were filtered using the goodGenes function with default values, and the filtered expression matrix was used for network construction. To determine an appropriate soft-thresholding power, we evaluated scale-free topology fit indices (pickSoftThreshold) across a range of powers (1–20), and a power of 11 was selected. The adjacency matrix was constructed using signed Pearson correlations and transformed into a topological overlap matrix (TOM). Genes were hierarchically clustered based on TOM dissimilarity, and initial modules were detected using dynamic tree cutting with minClusterSize = 100 and deepSplit = 2. Module colours were assigned, and eigengenes (first principal components of module expression) were calculated. Modules with highly similar eigengenes (r > 0.6, corresponding to a dissimilarity threshold of 0.40) were merged, resulting in 18 modules. The final module eigengenes were subjected to one-way ANOVA to assess differences across four experimental groups (stiff ancestral, soft ancestral, soft selected and stiff selected). Raw p-values were adjusted using the false discovery rate (FDR) method, and modules were ranked by adjusted significance. The top four modules (Black, Brown, Blue and Tan) with adjusted p-value < 0.05 were chosen for further analysis

### Single-sample gene set enrichment analysis (ssGSEA)

Single-sample Gene Set Enrichment Analysis (ssGSEA) was performed using the GSEApy Python package (gseapy.ssgsea function). Log₂-transformed and filtered gene expression data were used as input, with four types of gene sets: module genes identified in this study (Blue, Brown, Black, and Tan modules), MSigDB (Molecular Signatures Database) Hallmark, PID (Pathway Interaction Database), and KEGG (Kyoto Encyclopedia of Genes and Genomes) pathway gene sets. The analysis was run with sample_norm_method=’rank’ to compute normalized enrichment scores (NES) for each gene set across samples. The enrichment matrix was generated by pivoting the res2d output based on gene set terms and sample names. Differential enrichment between sample groups was assessed using Student’s t-test, considering gene sets with p < 0.05 as significantly enriched.

### Differential pathway analysis

A Student’s *t*-test was performed on the PID pathway enrichment scores obtained from ssGSEA to identify significantly enriched pathways across conditions. Results were visualized using a volcano plot, applying thresholds of *p* < 0.05 and |*t*-statistic| > 5. Principal Component Analysis (PCA) was also conducted on these enrichment scores following the same methodology described in the PCA section.

### Gene ontology

Gene Ontology (GO) enrichment analysis for module genes was performed using the GSEApy Python package. Genes from each co-expression module (Blue, Brown, Black, and Tan) were individually analysed for enrichment in GO Biological Process (GOBP 2025) pathways using the gp.enrichr() function. The analysis was conducted with the Human gene background, and all significantly enriched GO terms (adjusted p < 0.05) were retained. Enrichment results were stored and compared across modules to identify overrepresented biological processes associated with each gene module.

### Assay for Transposase-Accessible Chromatin with Sequencing (ATAC-seq)

Nuclei were isolated from 200,000 cells and ATAC-seq libraries were prepared on aliquots of 50,000 nuclei using an ATAC-seq library preparation kit (Active Motif). Pooled libraries were sequenced on a Novaseq 6000 S4 2×150 flow cell (ICBR Next-Gen Sequencing Core, University of Florida) at a depth of 100 million reads per sample. The data has been deposited under the GSE ID GSE255827.

### ATAC-Seq data analysis

Raw paired-end FASTQ files were trimmed for adapter sequences and low-quality bases with fastp, and read quality before and after trimming was assessed with FastQC. Trimmed reads were aligned to the indexed human reference genome GRCh38 (no-alt assembly) using bowtie2 with parameters --local -k 1 --no-mixed --no-discordant. Alignment files were converted to BAM, sorted, and indexed using samtools. Mitochondrial reads (chrM) were excluded, and PCR/optical duplicates were marked with Picard MarkDuplicates (with REMOVE_DUPLICATES=false and CREATE_INDEX=true). Low-quality and non-properly paired reads were filtered using samtools view with parameters -f 2 -F 1548 -q 30. To eliminate artifacts, reads overlapping ENCODE hg38 blacklist regions (v2) were removed using bedtools intersecting with the -v option. The resulting blacklist-filtered BAM files were sorted and indexed for downstream analysis. Normalized genome-wide coverage tracks were generated using bamCoverage (deepTools) with parameters --binSize 50 --normalizeUsing CPM --smoothLength 200 --effectiveGenomeSize 2913022398, producing CPM-normalized bigWig files. Peak calling was performed with MACS3 using the parameters -f BAMPE -q 0.01, generating peaks from the blacklist-filtered BAM files. Peaks were then mapped to gene promoters using a custom Python script with gene coordinates from the NCBI RefSeq annotation (Index of /refseq/H_sapiens/annotation/GRCh38_latest/refseq_identifiers). Only protein-coding genes were retained for downstream analysis. To quantify accessibility, bedtools multicov was used to calculate raw read counts across all merged peaks in each sample. Read counts were then aggregated per region, and DESeq2 was used to identify differentially accessible regions and estimate fold-changes. For visualization purposes, reads overlapping MACS3-called peak regions were retained to generate peak-restricted BAM files. CPM-normalized coverage tracks were regenerated from these peak-restricted BAM files using bamCoverage with the same parameters described above. The resulting bigWig files were visualized using the Integrative Genomics Viewer (IGV), ensuring that the displayed signal reflects chromatin accessibility specifically at confidently identified peak regions and minimizes background signal.

### Reduced Representation Bisulfite Sequencing (RRBS)

The RRBS-seq data in MDA-MB-231 ancestral and selected samples on soft (1 kPa) and stiff (308 kPa) was obtained from GSE255831.

### RRBS data analysis

The human reference genome (Homo_sapiens.GRCh38.dna.toplevel.fa.gz) was downloaded from Ensembl and prepared for bisulfite mapping using Bismark’s genome preparation module with the --bowtie2 option to create the bisulfite-converted indices. Raw FASTQ files from RRBS libraries were trimmed for adapter sequences and low-quality bases using Trim Galore with the --rrbs option. Trimmed reads were assessed for quality using FastQC, checking for read quality, adapter removal, and trimming efficiency. Trimmed reads were aligned to the prepared bisulfite-converted GRCh38 genome using Bismark (default parameters). Following alignment, Bismark methylation extractor was used to determine the CpG methylation levels from the BAM files. The resulting coverage files were used to estimate the differentially methylated CpG sites using the methylKit R package.

## Author contributions

Conceptualized and designed research: S.S., T.P.L. and M.K.J.; supervised research: T.P.L., and M.K.J.; performed research: S.S., S.K. T.W; interpreted data: S.S., S.K., T.W., I.S., T.P.L. and M.K.J.; funding acquisition: T.P.L., and M.K.J.; manuscript writing/editing: S.S. (prepared first draft), S.K., T.P.L., and M.K.J. (edited with inputs from all authors).

## Declaration of interests

The authors declare no conflicts of interest.

## Acknowledgments

S.S. is supported by PMRF (Prime Minister’s Research Fellowship) awarded by the Government of India. M.K.J. is supported by Param Hansa Philanthropies. T.P.L. is supported by the National Institutes of Health (NIH) grant U01 CA225566 and the Cancer Prevention and Research Institute of Texas (CPRIT) Established Investigator Award RR200043. We are grateful to Dr. Irtisha Singh for her suggestions. We also thank Ms. Kajal Charan and Dr. Paras Jain for their valuable insights concerning the development of the computational models used in this study.

## Supplementary Figures

**Supplementary Figure S1:**
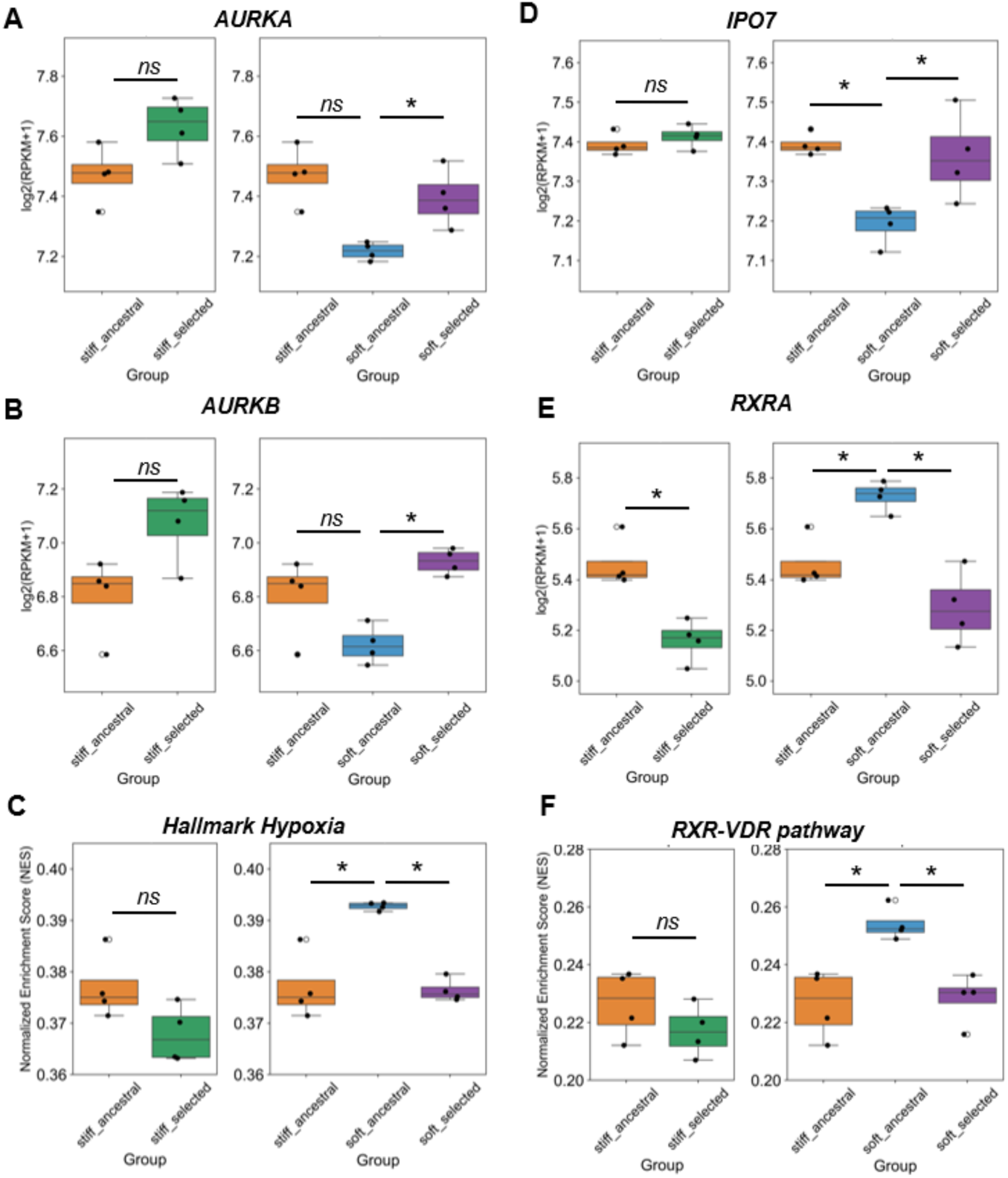
Gene expression values for individual genes or ssGSEA scores for pathways for **(A)** AURKA gene; **(B)** AURKB gene; **(C)** MSigDB Hallmark Hypoxia pathway; **(D)** IPO7 gene; **(E)** RXRA gene and **(F)** PID RXR-VDR pathways across the two different trajectories. Asterisks indicate statistical significance based on Students t-test with p-value < 0.05

**Supplementary Figure S2:**
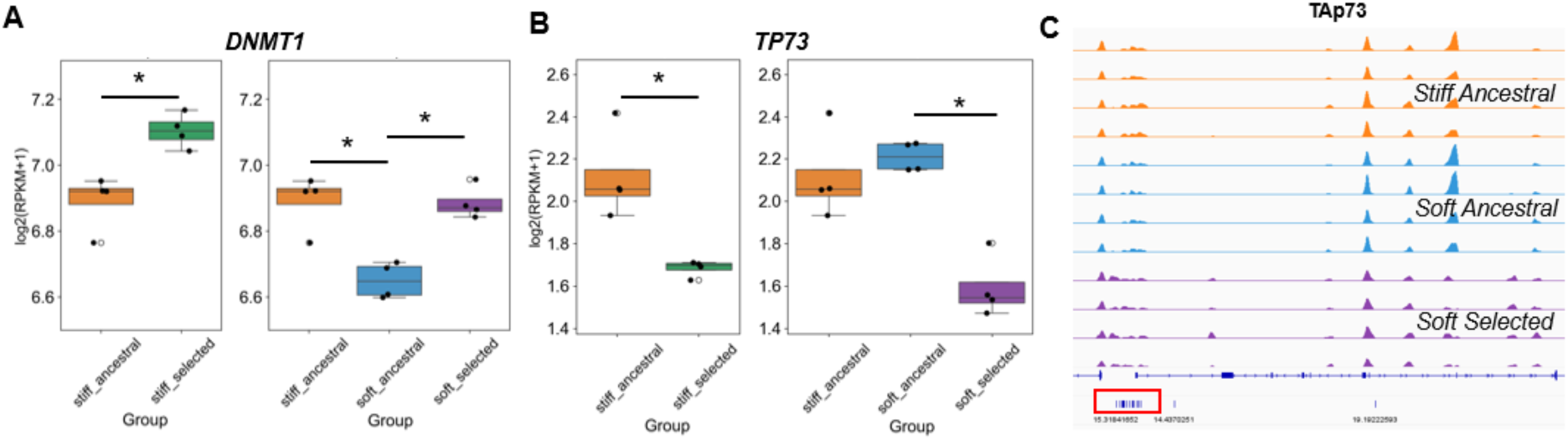
Gene expression values for individual genes (A) DNMT1 and (B) TAp73. (C) Genome browser tracks showing ATAC-seq (accessibility) and RRBS (methylation) data at the TAp73 promoter for four replicates each of stiff ancestral (orange), soft ancestral (blue) and soft selected (purple).

**Supplementary Figure S3:**
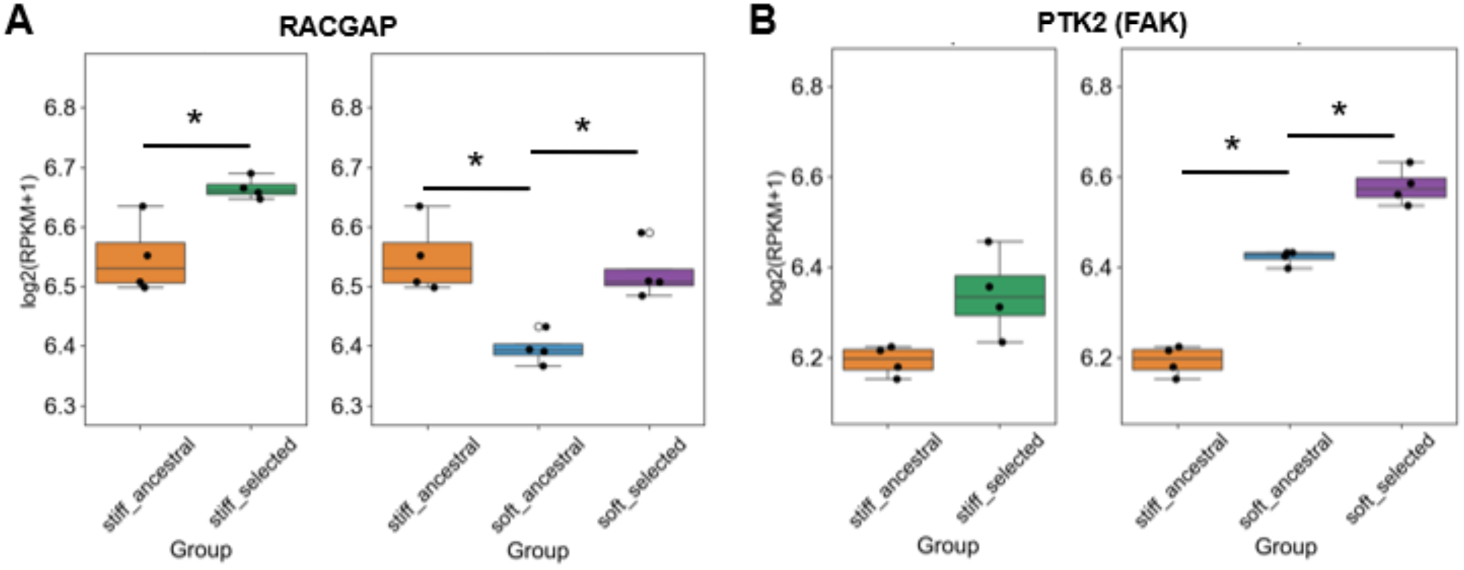
Gene expression values for individual genes (A) RACGAP1 and (B) PTK2 (FAK).

## Notes

### Competing Interest Statement

The authors have declared no competing interest.

