## Supplementary Information for "Multi-omics investigation reveals molecular determinants of cancer cell evolution on soft extracellular matrix"

### Module 1: YAP/TAZ Mechanotransduction

Module 1 is adapted from previously published mechanotransduction models that describe the canonical stiffness-sensing pathway involving FAK activation, RhoA cycling, ROCK and mDia signaling, Myosin contractility, LIMK–Cofilin–actin turnover, and cytoplasmic YAP/TAZ phosphorylation [1]. These components form the established mechanochemical cascade linking matrix stiffness to cytoskeletal tension and YAP/TAZ activation. In this work, the model is extended by incorporating additional layers of regulation that were not present in earlier formulations, including explicit equations for nuclear YAP/TAZ dynamics, Importin-dependent nuclear transport, NPC activation governed by LaminA and cytoskeletal tension, a stress-fiber remodeling term sensitive to substrate stiffness, and transcriptional feedback through YAP-dependent Importin and MYBL2 production along with YAP–MYBL2 complex formation.

#### FAK and pFAK Dynamics

In the FAK/pFAK module, FAK is phosphorylated to its active form pFAK through the rate $R_1 = k_f\, FAK + k_{sf} \frac{E}{C+E} FAK$, where $k_f$ denotes basal phosphorylation and $k_{sf} \frac{E}{C+E}$ represents stiffness-dependent activation that increases with extracellular stiffness $E$. pFAK is dephosphorylated back to FAK at the rate $R_2 = k_{df}\, pFAK$. In addition to this fast phosphorylation cycle, total FAK levels adjust more slowly through a production term $k_{\mathrm{FAK}}^{\mathrm{prod}} \left(1 - \frac{FAK+pFAK}{K_{\mathrm{FAK}}}\right) SF_{\mathrm{remod}}$, which promotes FAK synthesis during stress-fiber remodeling, and a degradation term $k_{\mathrm{FAK}}^{\mathrm{deg}} FAK\, SF_{\mathrm{remod}}$. Together, these reactions describe how mechanical inputs rapidly regulate pFAK levels while cytoskeletal remodeling tunes the overall FAK abundance over longer timescales.

$$\frac{dFAK}{dt} = R_2 - R_1 + k_{\text{FAK}}^{\text{prod}} \left(1 - \frac{FAK + pFAK}{K_{\text{FAK}}}\right) SF_{\text{remod}} - k_{\text{FAK}}^{\text{deg}}\, FAK\, SF_{\text{remod}}$$

$$\frac{dpFAK}{dt} = R_1 - R_2$$

where

$$R_1 = k_f FAK + k_{sf} \frac{E}{C+E} FAK, \qquad R_2 = k_{df}\, pFAK.$$

#### RhoA Dynamics

RhoA cycles between its inactive GDP-bound form ($RhoA_{GDP}$) and active GTP-bound form ($RhoA_{GTP}$). Activation occurs through

$$R_3 = (k_{fk\rho}(\gamma\, pFAK^n + 1) + k_{\rho,mybl2}\, yap\_mybl2)\, RhoA_{GDP},$$

where the term $k_{fk\rho}(\gamma pFAK^n + 1)$ represents FAK-driven activation, reflecting how increasing pFAK enhances RhoA activation. The term $k_{\rho,mybl2}\, yap\_mybl2$ introduces MYBL2-mediated transcriptional feedback via the YAP-MYBL2 complex. RhoA deactivation is governed by

$$R_4 = k_{d\rho}\, RhoA_{GTP},$$

representing GAP-mediated conversion of RhoA–GTP back to the GDP-bound state. The overall dynamics are given by

$$\frac{dRhoA_{GDP}}{dt} = -R_3 + R_4, \qquad \frac{dRhoA_{GTP}}{dt} = R_3 - R_4,$$

ensuring mass conservation between the two forms. This formulation captures the integration of mechanical signals via pFAK and transcriptional reinforcement through MYBL2 in tuning RhoA activation.

#### ROCK Dynamics

ROCK transitions between an inactive form ($ROCK$) and its active form ($ROCK_A$). Deactivation occurs at

$$R_5 = k_{drock}\, ROCK_A,$$

which represents the intrinsic deactivation rate of active ROCK. Activation is driven by RhoA–GTP through

$$R_6 = k_{r\rho}\, RhoA_{GTP}\, ROCK,$$

indicating that greater levels of active RhoA promote ROCK activation in proportion to the available inactive ROCK pool. The resulting dynamics are

$$\frac{dROCK}{dt} = R_5 - R_6, \qquad \frac{dROCK_A}{dt} = -R_5 + R_6.$$

These equations represent the central role of RhoA-GTP in activating ROCK, which subsequently regulates downstream cytoskeletal tension, modulation of Myosin activity, and actin remodeling via LIMK–Cofilin signaling.

#### mDia Dynamics

mDia cycles between an inactive form ($mDia$) and an active form ($mDia_A$). Deactivation of active mDia occurs through

$$R_7 = k_{dmdia}\, mDia_A,$$

representing its intrinsic inactivation rate. Activation is driven by RhoA–GTP according to

$$R_8 = k_{m\rho}\, RhoA_{GTP}\, mDia,$$

showing that RhoA–GTP promotes mDia activation proportionally to the pool of inactive mDia. The governing equations are

$$\frac{dmDia}{dt} = R_7 - R_8, \qquad \frac{dmDia_A}{dt} = -R_7 + R_8.$$

Overall, it captures how RhoA signaling activates mDia, a formin protein that nucleates and elongates actin filaments, shaping cytoskeletal organization and contributing to force generation and mechanotransduction.

#### Myosin Dynamics

Myosin activity is regulated by ROCK-dependent phosphorylation. To capture the non-linear dependency on ROCK activity, we define a thresholded active ROCK term, $T_{ROCKA}$:

$$T_{ROCKA} = \frac{1}{2}\left[\tanh\left(sc_1(ROCK_A - ROCK_b)\right) + 1\right] ROCK_A,$$

where $sc_1$ controls the steepness and $ROCK_b$ is the activation threshold. Myosin activation occurs at rate $R_9$, driven by this thresholded ROCK term:

$$R_9 = k_{mr}(\epsilon T_{ROCKA} + 1)Myo - k_{dmy} Myo_A.$$

Here, $Myo$ converts to active myosin ($Myo_A$, representing phosphorylated myosin light chain) based on ROCK activity scaled by $\epsilon$. The term $+1$ represents a basal activation rate. Deactivation occurs at a constant rate $k_{dmy}$. The dynamics are:

$$\frac{dMyo}{dt} = -R_9, \qquad \frac{dMyo_A}{dt} = R_9.$$

#### LIMK Dynamics

LIMK (LIM kinase) is another direct downstream effector of ROCK, responsible for inhibiting Cofilin. Its activation kinetics follow a similar ROCK-dependency as Myosin:

$$R_{10} = k_{lr}(\tau T_{ROCKA} + 1)LIMK - k_{dl} LIMK_A.$$

Active ROCK promotes the transition of inactive $LIMK$ to active $LIMK_A$, scaled by parameter $\tau$. Deactivation follows first-order kinetics with rate $k_{dl}$.

$$\frac{dLIMK}{dt} = -R_{10}, \qquad \frac{dLIMK_A}{dt} = R_{10}.$$

#### Cofilin Dynamics

Cofilin acts as an actin-severing protein, promoting depolymerization. It is inactivated via phosphorylation by LIMK. The reaction rate $R_{11}$ represents the net flux between the phosphorylated (inactive) and non-phosphorylated (active) states:

$$R_{11} = k_{turn\_over} Cofilin_p - \frac{k_{catcofilin} \cdot LIMK_A \cdot Cofilin_{NP}}{k_{mcofilin} + Cofilin_{NP}}.$$

The first term, $k_{turn\_over} Cofilin_p$, represents the reactivation (dephosphorylation) of Cofilin. The second term is the phosphorylation of active Cofilin ($Cofilin_{NP}$) by LIMK, modeled using Michaelis-Menten kinetics. The differential equations are:

$$\frac{dCofilin_p}{dt} = -R_{11}, \qquad \frac{dCofilin_{NP}}{dt} = R_{11}.$$

Thus, when LIMK is active, $R_{11}$ becomes negative, shifting the balance toward inactive $Cofilin_p$, thereby stabilizing F-actin.

#### Actin Dynamics

The interconversion between globular actin ($G\_actin$) and filamentous actin ($F\_actin$) determines cytoskeletal integrity. Polymerization is promoted by ROCK (via downstream effectors like mDia and inhibition of depolymerization), while depolymerization is facilitated by active Cofilin.

$$R_{12} = k_{ra}(\alpha T_{ROCKA} + 1) G\_actin - (k_{dep} + k_{fc1} Cofilin_{NP}) F\_actin.$$

The first term represents polymerization enhanced by active ROCK (scaled by $\alpha$). The second term represents depolymerization, which occurs at a basal rate $k_{dep}$ and is significantly accelerated by active Cofilin ($k_{fc1} Cofilin_{NP}$).

$$\frac{dG\_actin}{dt} = -R_{12}, \qquad \frac{dF\_actin}{dt} = R_{12}.$$

#### Cytosolic YAP/TAZ Phosphorylation

The phosphorylation status of YAP/TAZ regulates its subcellular localization. Phosphorylated YAP ($YAPTAZ_p$) is sequestered in the cytoplasm or degraded, while unphosphorylated YAP ($YAPTAZ$) is capable of nuclear entry. Mechanical tension inhibits the phosphorylation complex (e.g., LATS), effectively dephosphorylating YAP.

$$R_{13} = (k_{CN} + k_{CY} \cdot F\_actin \cdot Myo_A) YAPTAZ_p - k_{NC} YAPTAZ.$$

Here, $k_{CN}$ is the basal dephosphorylation rate, and $k_{CY} \cdot F\_actin \cdot Myo_A$ represents tension-dependent dephosphorylation derived from actomyosin contractility. The term $k_{NC}$ represents the basal phosphorylation rate (kinase activity). The mass balance for the cytosolic pools is:

$$\frac{dYAPTAZ_p}{dt} = -R_{13}, \qquad \frac{dYAPTAZ}{dt} = R_{13} - R_{16},$$

where $R_{16}$ (defined later) is the flux of unphosphorylated YAP into the nucleus.

#### Lamin A and NPC Activation

Nuclear cytoskeletal coupling modulates the permeability of the nuclear envelope. First, we define the cytosolic stiffness stress $E_{cytosol}$ exerted by the actin network:

$$E_{cytosol} = p \cdot (F\_actin)^{2.6}.$$

Lamin A phosphorylation is modeled as a response to this stress, softening the lamina:

$$R_{14} = k_{fl} \frac{E_{cytosol}}{C_{Lamin} + E_{cytosol}} LaminA_p - k_{rl} LaminA.$$

Consequently, Nuclear Pore Complexes (NPCs) are mechanically activated (opened) by the combined action of Lamin A and actomyosin tension:

$$R_{15} = k_{fNPC} \cdot LaminA \cdot F\_actin \cdot Myo_A \cdot NPC - k_{rNPCA} NPC_A.$$

This implies that NPC opening ($NPC_A$) requires structural support from Lamin A and pulling forces from the cytoskeleton.

$$\frac{dNPC}{dt} = -R_{15}, \qquad \frac{dNPC_A}{dt} = R_{15}.$$

#### YAP/TAZ Nuclear Transport

The transport of unphosphorylated YAP into the nucleus depends on both the availability of transporters (Importin) and the mechanical state of the nuclear pores (NPCs).

$$R_{16} = \left( k_{inb\_max} \frac{Importin}{3} + k_{in} NPC_A \right) YAPTAZ - k_{out} YAPTAZ_{nuc}.$$

The import rate consists of a basal term dependent on Importin concentration and a mechanically-regulated term dependent on active NPCs. Export occurs at a constant rate $k_{out}$.

#### Transcriptional Feedback and MYBL2 Complex

Nuclear YAP ($YAPTAZ_{nuc}$) acts as a transcriptional co-activator for several targets, including Importin (specifically IPO7) and MYBL2.

**Importin Production:**

$$\frac{dImportin}{dt} = basal_{imp} + k_{prod}^{imp} \frac{YAPTAZ_{nuc}^{h_{imp}}}{K_{imp}^{h_{imp}} + YAPTAZ_{nuc}^{h_{imp}}} - k_{deg}^{imp} Importin.$$

This Hill function models upregulation of nuclear transport machinery by YAP, creating a positive feed-forward loop.

**MYBL2 Production and Complex Formation:** Similarly, MYBL2 production is driven by nuclear YAP:

$$NetProd_{mybl2} = basal_{mybl2} + k_{prod}^{mybl2} \frac{YAPTAZ_{nuc}^{h_{mybl2}}}{K_{mybl2}^{h_{mybl2}} + YAPTAZ_{nuc}^{h_{mybl2}}} - k_{deg}^{mybl2} mybl2.$$

Within the nucleus, MYBL2 interacts with YAP to form a transcriptional complex ($yap\_mybl2$). The rate of complex formation is:

$$\frac{d(yap\_mybl2)}{dt} = k_{comp} \cdot mybl2 \cdot YAPTAZ_{nuc} - k_{diss} \cdot yap\_mybl2.$$

To conserve mass, the derivatives for free nuclear YAP and free MYBL2 account for this complex formation:

$$\frac{dYAPTAZ_{nuc}}{dt} = R_{16} - \frac{d(yap\_mybl2)}{dt},$$
$$\frac{dmybl2}{dt} = NetProd_{mybl2} - \frac{d(yap\_mybl2)}{dt}.$$

##### Stress Fiber Remodeling

Finally, the model accounts for the long-term remodeling of stress fibers ($SF_{remod}$) based on substrate stiffness $E$. Remodeling occurs only when stiffness is below a silencing threshold ($E_{sil\_thresh}$):

$$\frac{dSF_{remod}}{dt} = k^f_{remod} F\_actin - k^d_{remod} SF_{remod},$$

where $k^f_{remod}$ and $k^d_{remod}$ are effective rates that become zero if $E \geq E_{sil\_thresh}$. This variable feeds back into the slow production of FAK (described in the FAK section), allowing the cell to adapt its sensitivity based on exposure to chronic stiffness.

#### Module 2: Cell Cycle Regulation

Adapted from the previously published work [2], this module describes the core cell cycle oscillator, centered on the interaction between Cyclin B/Cdk1 and the Anaphase-Promoting Complex/Cyclosome (APC/C) co-activators Cdc20 and Cdh1. The dynamics are calculated in hours ($h^{-1}$). To account for global physiological scaling or non-dimensionalization present in the computational implementation, most derivatives are scaled by a factor $d$.

Crucially, this module is coupled to Module 1 via two specific mechanisms:

1. **Transcriptional Activation:** The YAP-MYBL2 complex ($yap\_mybl2$) from Module 1 promotes the transcription of Cdc20.

2. **Epigenetic Silencing:** The stress fiber remodeling variable ($SF_{remod}$) drives an epigenetic silencing factor ($EpiSil$), which represses Cdh1 mRNA production.

##### Cyclin B Dynamics

Cyclin B drives mitotic entry. Its messenger RNA ($B_m$) is produced in response to Growth Factors ($GF$) following saturation kinetics and degraded linearly. The protein ($CycB$) is synthesized from $B_m$. It undergoes basal degradation ($k_{2a}$) and ubiquitin-mediated degradation catalyzed by active Cdh1 ($k_{2b}$).

$$\frac{dB_m}{dt} = d \cdot \left( \frac{k_{1m} \cdot GF}{k_{mm} + k_{eff} \cdot GF} - k_{1dm} B_m \right)$$

$$\frac{dCycB}{dt} = d \cdot (k_1 B_m - k_{2a} CycB - k_{2b} CycB \cdot Cdh1)$$

#### Cdc20 Dynamics and Mechanotransduction Link

Cdc20 is required for the metaphase-to-anaphase transition. Its mRNA ($C20_m$) production is regulated by a positive feedback loop from Cyclin B and, critically, by the YAP-MYBL2 complex from Module 1.

We define the activation term from Module 1 as a Hill function of the nuclear complex:

$$\Psi_{mybl2} = \frac{(yap\_mybl2)^{h_{mybl2\_cdc}}}{K_{mybl2\_cdc}^{h_{mybl2\_cdc}} + (yap\_mybl2)^{h_{mybl2\_cdc}}}$$

The influence of Cyclin B is also modeled as a Hill function:

$$H_{cycb} = \frac{CycB^{n_{cc}}}{J_5^{n_{cc}} + CycB^{n_{cc}}}$$

The total production rate of Cdc20 mRNA is the product of these regulatory inputs. The protein exists in a total pool ($Cdc20_{tot}$) and an active form ($Cdc20_A$).

$$\frac{dC20_m}{dt} = d \cdot \left[ \left( k_{5am} + \frac{k_{5bm} H_{cycb}}{k_{5cm} + GF \cdot j_{5c}} \right) \Psi_{mybl2} - k_{5dm} C20_m \right]$$

$$\frac{dCdc20_{tot}}{dt} = d \cdot (k_{5a} C20_m - k_6 Cdc20_{tot})$$

Active Cdc20 ($Cdc20_A$) is generated from the inactive pool ($Cdc20_{tot} - Cdc20_A$) by the Intermediary Enzyme (IEP) and inhibited by the Spindle Assembly Checkpoint (represented by $Mad$).

$$\frac{dCdc20_A}{dt} = d \cdot \left( \frac{k_7 \cdot IEP \cdot (Cdc20_{tot} - Cdc20_A)}{J_7 + (Cdc20_{tot} - Cdc20_A)} - \frac{k_8 \cdot Mad \cdot Cdc20_A}{J_8 + Cdc20_A} - k_6 Cdc20_A \right)$$

#### Cdh1 Dynamics and Epigenetic Silencing

Cdh1 keeps the cell in G1 phase by degrading Cyclins. Its expression is sensitive to matrix stiffness via an epigenetic silencing mechanism.

The silencing factor ($S_{factor}$) reduces Cdh1 mRNA ($cdh_m$) production based on the level of $EpiSil$. Furthermore, the basal production rate $k_{3m}$ increases effectively to $k_{3m}^{eff}$ (by factor 1.73) when stiffness $E$ is below a threshold $E_{sil\_thresh}$, simulating contact inhibition or low-stiffness quiescence.

$$S_{factor} = \frac{1}{1 + k_{sil\_strength} \cdot EpiSil}$$

$$\frac{dcdh_m}{dt} = d \cdot \left( k_{3m}^{eff} \cdot S_{factor} - k_{3dm} cdh_m \right)$$

$$\frac{dcdh_{tot}}{dt} = d \cdot (k_{3a} cdh_m - k_{3dt} cdh_{tot})$$

Active Cdh1 is regulated by phosphorylation. It is activated by Cdc20A and deactivated by Cyclin B (Cdk1).

$$\frac{dCdh1}{dt} = d \cdot \left( \frac{(k_3 + k_{3b} Cdc20_A)(cdh_{tot} - Cdh1)}{J_3 + (cdh_{tot} - Cdh1)} - \frac{k_4 \cdot CycB \cdot Cdh1}{J_4 + Cdh1} - k_{3dt} Cdh1 \right)$$

#### IEP and Cdt1 Dynamics

The Intermediary Enzyme (IEP) acts as a trigger for Cdc20 activation, driven by Cyclin B. Cdt1, a replication licensing factor, is degraded by Cyclin B, ensuring DNA replication occurs only once per cycle.

$$\frac{dIEP}{dt} = d \cdot (k_9 \cdot CycB \cdot (1 - IEP) - k_{10} IEP)$$

$$\frac{dcdt1}{dt} = d \cdot (k_{11} - k_{12} \cdot CycB \cdot cdt1 - k_{13} cdt1)$$

#### Epigenetic Silencing Variable

The variable $EpiSil$ represents the accumulation of epigenetic marks (e.g., DNA methylation) that silence tumor suppressors like Cdh1. This accumulation is driven by the stress fiber remodeling variable ($SF_{remod}$) from Module 1 via a steep Hill function, linking chronic mechanical stress to long-term cell cycle deregulation.

$$\frac{dEpiSil}{dt} = k_{epi}^{on} \frac{SF_{remod}^3}{(Th_{remod})^3 + SF_{remod}^3} - k_{deg}^{epi} EpiSil$$

Note: Unlike other cell cycle variables, the $EpiSil$ dynamic is not scaled by $d$ in this implementation, reflecting its operation on a distinct timescale.

### Computational Modeling of Clonal Selection Dynamics

To simulate the evolutionary selection of cellular subpopulations under resource-limited conditions, we developed a discrete-time stochastic model using Python3.

**Data Pre-processing and Initialization** Input proliferation data (`avg_divisions_phase2`) was filtered to exclude non-viable clones (rate $\leq 0$) and randomized to ensure unbiased initial ordering. Division rates were normalized to a daily time scale (derived from 10-day aggregate intervals). The simulation was initialized with a heterogeneous population of $N$ clones, where $N$ represents the number of unique division rates identified. Each clone was assigned an equal initial population size of 5 arbitrary units.

**Simulation Parameters** The model was executed over 90 time steps, representing a 90-day period. For each clone $i$, the net growth rate ($R_{\text{net},i}$) was calculated by incorporating a fixed cell death penalty to account for turnover. The death rate was set at 20% of the birth rate, yielding:

$$R_{\text{net},i} = R_{\text{birth},i} \times (1 - 0.20) \tag{1}$$

**Selection Algorithm** At each time step $t$, the population dynamics were governed by a two-phase update cycle:

1. **Growth Phase:** The population of each clone was updated based on its specific net growth rate according to:

$$P_{i,t+1} = P_{i,t} \times (1 + R_{\text{net},i}) \tag{2}$$

2. **Competition Phase:** To model spatial or nutrient constraints (Carrying Capacity), the total population was normalized at the end of every time step. The aggregate population was forced to return to the initial total population count ($P_{\text{total},t=0}$), thereby scaling down all clones proportionally. This step enforced a zero-sum competition where an increase in the frequency of faster-cycling clones necessitated a decrease in slower-cycling clones.

**Quantification of Evolutionary Fitness** To visualize the trajectory of population adaptation, we calculated the weighted average division rate of the entire population at each time step. This metric served as a proxy for the overall evolutionary fitness of the tumor, defined as:

$$\text{Fitness}_{\text{avg}}(t) = \frac{\sum (P_{i,t} \times R_{\text{net},i})}{\sum P_{i,t}} \tag{3}$$

Results were visualized using stacked bar charts to display clonal composition changes and line plots to track the shift in global fitness over time.

### Supplementary Tables: Model Parameters

Table 1: **Module 1: YAP/TAZ Mechanotransduction Parameters**

| **Symbol** | **Description** | **Value** |
|---|---|---|
| $k_f$ | Basal FAK activation rate | 54.0 |
| $k_{sf}$ | Max stiffness-dependent FAK activation | 1364.4 |
| $k_{df}$ | FAK deactivation rate | 126.0 |
| $C$ | Stiffness half-maximal constant (FAK) | 3.25 |
| $k_{FAK}^{prod}$ | FAK production rate | 1260.0 |
| $k_{deg}^{FAK}$ | FAK degradation rate | 540.0 |
| $K_{FAK}$ | FAK upregulation constant | 1.0 |
| $k_{fk\_rho}$ | RhoA activation rate by FAK | 60.48 |
| $\gamma$ | FAK-dependent gain on RhoA | 77.56 |
| $n$ | Hill coefficient for RhoA activation | 5 |
| $k_{d\rho}$ | RhoA deactivation rate | 2250.0 |
| $k_{\rho,mybl2}$ | RhoA activation by MYBL2 complex | 5.4 |
| $k_{r\rho}$ | ROCK activation by RhoA | 2332.8 |
| $k_{drock}$ | ROCK deactivation rate | 2880.0 |
| $k_{m\rho}$ | mDia activation by RhoA | 7.2 |
| $k_{dmdia}$ | mDia deactivation rate | 18.0 |
| $sc_1$ | ROCK threshold steepness | 20 |

Continued on next page...

**Table 1 – continued from previous page**

| Symbol | Description | Value |
|---|---|---|
| $ROCK_b$ | ROCK activation threshold | 0.3 |
| $k_{mr}$ | Myosin activation by ROCK | 108.0 |
| $\epsilon$ | ROCK-dependent gain on Myosin | 36 |
| $k_{dmy}$ | Myosin deactivation rate | 241.2 |
| $k_{lr}$ | LIMK activation by ROCK | 252.0 |
| $\tau$ | ROCK-dependent gain on LIMK | 55.49 |
| $k_{dl}$ | LIMK deactivation rate | 7200.0 |
| $k_{turn\_over}$ | Cofilin reactivation (dephosphorylation) | 144.0 |
| $k_{catcofilin}$ | Cofilin phosphorylation rate | 1224.0 |
| $k_{mcofilin}$ | Cofilin Michaelis constant | 14400.0 |
| $k_{ra}$ | Actin polymerization rate | 1440.0 |
| $\alpha$ | ROCK-dependent gain on polymerization | 50 |
| $k_{dep}$ | Basal actin depolymerization rate | 12600.0 |
| $k_{fc1}$ | Cofilin-dependent depolymerization rate | 14400.0 |
| $p$ | Cytosolic stiffness coefficient | $9 \times 10^{-6}$ |
| $k_{CN}$ | Basal YAP dephosphorylation rate | 2016.0 |
| $k_{CY}$ | Tension-dependent YAP dephosphorylation | 2.736 |
| $k_{NC}$ | Basal YAP phosphorylation rate | 504.0 |
| $k_{in}$ | NPC-dependent YAP import rate | 36000.0 |
| $k_{inb\_max}$ | Basal (Importin-dependent) YAP import | 3600.0 |
| $k_{out}$ | YAP export rate | 3600.0 |
| $k_{fl}$ | Lamin phosphorylation rate | 1656.0 |
| $C_{Lamin}$ | Lamin stiffness half-max constant | 100 |
| $k_{rl}$ | Lamin dephosphorylation rate | 3.6 |
| $k_{fNPC}$ | NPC activation rate | $1.0 \times 10^{-3}$ |
| $k_{rNPCA}$ | NPC deactivation rate | 31320.0 |
| $k_{prod}^{imp}$ | Inducible Importin production | 1404.0 |
| $basal_{imp}$ | Basal Importin production | 9396.0 |
| $k_{deg}^{imp}$ | Importin degradation | 3600.0 |
| $K_{imp}$ | Importin Hill constant | 0.4 |
| $h_{imp}$ | Importin Hill coefficient | 4 |
| $k_{prod}^{mybl2}$ | Inducible MYBL2 production | 720.0 |

Continued on next page...

**Table 1 – continued from previous page**

| **Symbol** | **Description** | **Value** |
|---|---|---|
| $basal_{mybl2}$ | Basal MYBL2 production | 36.0 |
| $k_{deg}^{mybl2}$ | MYBL2 degradation | 360.0 |
| $K_{mybl2}$ | MYBL2 Hill constant | 0.4 |
| $h_{mybl2}$ | MYBL2 Hill coefficient | 4 |
| $comp_{f\_ym}$ | YAP-MYBL2 complex formation | 1080.0 |
| $k_{diss\_ym}$ | YAP-MYBL2 complex dissociation | 1908.0 |
| $k_{remod}^{f}$ | Stress fiber remodeling (forward) | $3.6 \times 10^{-4}$ |
| $k_{remod}^{d}$ | Stress fiber remodeling (reverse) | 0.036 |
| $Th_{remod}$ | Threshold for epigenetic activation | 0.5 |

Table 2: **Module 2: Cell Cycle and Interface Parameters**

| **Symbol** | **Description** | **Value** |
|---|---|---|
| $d$ | Time scaling factor (physiological rate) | 2.8 |
| $GF$ | Growth Factor concentration | 2.0 |
| $k_{mm}$ | Michaelis constant for GF | 0.20 |
| $k_{eff}$ | GF efficacy constant | 1.0 |
| $k_{1m}$ | Basal Cyclin B mRNA production | 0.0037 |
| $k_{1dm}$ | Cyclin B mRNA degradation | 0.058 |
| $k_1$ | Cyclin B protein synthesis | 0.4 |
| $k_{2a}$ | Basal Cyclin B degradation | 0.04 |
| $k_{2b}$ | Cdh1-dependent Cyclin B degradation | 2.0 |
| $k_3$ | Basal Cdh1 activation | 1.28 |
| $k_{3a}$ | Cdh1 total protein synthesis | 1.0 |
| $k_{3b}$ | Cdc20A-dependent Cdh1 activation | 8.0 |
| $k_4$ | Cyclin B-dependent Cdh1 inactivation | 40.0 |
| $J_3, J_4$ | Michaelis constants for Cdh1 | 0.04 |
| $k_{3m}$ | Basal Cdh1 mRNA production | 0.5 |
| $k_{3dm}$ | Cdh1 mRNA degradation | 0.5 |
| $k_{3dt}$ | Cdh1 protein turnover | 1.0 |
| $k_{5am}$ | Basal Cdc20 mRNA production | 0.005 |
| $k_{5bm}$ | Inducible Cdc20 mRNA production | 0.2 |

Continued on next page...

**Table 2 – continued from previous page**

| Symbol | Description | Value |
|---|---|---|
| $k_{5cm}$ | Cdc20 mRNA inhibition constant | 1.0 |
| $j_{5c}$ | GF inhibition scaling | 0.02 |
| $k_{5dm}$ | Cdc20 mRNA degradation | 1.386 |
| $J_5$ | Hill constant (CycB effect on Cdc20) | 0.3 |
| $n_{cc}$ | Hill coefficient (CycB effect) | 4 |
| $k_{5a}$ | Cdc20 protein synthesis | 1.0 |
| $k_6$ | Cdc20 protein degradation | 0.05 |
| $k_7$ | IEP-dependent Cdc20 activation | 1.4 |
| $k_8$ | Mad-dependent Cdc20 inactivation | 0.5 |
| $J_7, J_8$ | Michaelis constants for Cdc20 | 0.001 |
| $Mad$ | Spindle Assembly Checkpoint signal | 1.0 |
| $k_9$ | IEP activation rate | 0.1 |
| $k_{10}$ | IEP inactivation rate | 0.02 |
| $k_{11}$ | Cdt1 synthesis rate | 0.045 |
| $k_{12}$ | Cyclin B-dependent Cdt1 degradation | 2.27 |
| $k_{13}$ | Basal Cdt1 degradation | 0.004 |
| $k_{epi}^{on}$ | Epigenetic silencing on-rate | 0.075 |
| $Epi_{deg}$ | Epigenetic silencing off-rate | 0.5 |
| $k_{sil\_strength}$ | Strength of epigenetic silencing on Cdh1 | 15 |
| $K_{mybl2\_cdc}$ | Hill constant (MYBL2 effect on Cdc20) | 0.172 |
| $h_{mybl2\_cdc}$ | Hill coefficient (MYBL2 effect) | 4 |
| $E_{sil\_thresh}$ | Stiffness threshold for remodeling | 1.5 |
